# Co-existence of Modularity and Anti-modularity in the Functional brain connectomes

**DOI:** 10.64898/2026.06.17.733035

**Authors:** Sheksha Dudekula, Aradhana Singh

**Affiliations:** Department of Physics, Indian Institute of Science Education and Research Tirupati, Tirupati 517619, Andhra Pradesh, India

**Keywords:** Functional connectome, Network Neuroscience, Modular Networks, Anti-modular Networks, k-Core, Complex Systems

## Abstract

The brain requires coordination among different regions to execute cognitive tasks, which may involve both positive- and negative-correlations. The topology of these correlations may indicate the mechanism underlying brain functioning in a given state. Here, we study changes in the functional connectomes (FCs) of both the positive and negative-correlations across various cognitive task states relative to the resting state, using publicly available electroencephalographic (EEG) data. Considering the EEG-specific topographical cortical regions as topographical modules (TMs), we find that the FC comprising positive correlations (*G*_+_) is modular. In contrast, networks of negative-correlations (*G*_*−*_) are anti-modular, with more connections between TMs than within them, and are associated with improved overall topological efficiency. These functional networks also show variability across frequency bands and brain states. In the low-frequency delta band, resting states exhibit higher modularity and anti-modularity than task states; in contrast, in the high-frequency Gamma band, modularity and anti-modularity are much higher during task states than in the resting state. The k-core analysis of all networks further reveals differences: *G*_+_ is more hierarchical and robust than *G*_*−*_ across all states. Moreover, the task-state networks are always more hierarchical than the resting-state networks across all frequency bands. In the high-frequency gamma band, they are also significantly more robust than the resting-state networks. These networks also differ in the topology of their innermost core constituents: the innermost core regions of *G*_+_ are randomly connected and spatially localized, mostly in posterior brain regions across subjects, in the high-frequency gamma band. Whereas those in *G*_*−*_ are spatially de-localized, cover the extreme anterior and extreme posterior brain regions, and remain anti-modular in all the frequency bands. Overall, our analysis reveals the presence of an anti-modular organization of functionally specialized TMs alongside their modular organization and points to task- and resting-state differences in their topologies.

## 1 Introduction

The orchestration of activities across multiple brain regions can provide an estimate of the extent to which they collaborate during a task. For example, coarsely speaking, crossing the road involves looking around for safe conditions and moving simultaneously, indicating potential cross-talk between the brain regions responsible for visual perception and motor movement control. Moreover, in terms of brain region activity, there can be many states, such as resting, listening, dancing, focusing, performing tasks, and solving problems. Functional brain networks, also called the functional connectome (FC), constructed from correlations or coherence in brain activity across different brain regions, provide an effective approach for understanding the brain in action [1]. The FC corresponding to each state reflects the underlying brain mechanism involved in understanding and processing during that state and can therefore reveal these mechanisms across different states. Further, in understanding FCs, network theory serves as a powerful tool. Network-based analysis has provided useful insights into how the brain ages and how a pathological condition affects it [2–5]. Moreover, it sheds light on how the particular cognition affects overall brain dynamics, such as how listening to music alters FC connectivity, and enhances the small-worldness in the presence of background noise compared to the silent background [6–8]. Small-world topology is associated with higher efficiency, as reflected in the higher clustering observed in most real-world networks [9, 10]. Two of the primary techniques for recording brain activity are functional magnetic resonance imaging (fMRI), which measures hemodynamic activity across different brain regions, and the electroencephalogram (EEG), which records electrical potentials from different brain regions. These recorded electrical potentials, often called brain waves, reflect the voltage shifts caused by the simultaneous firing of numerous neurons. The brain waves span many frequency bands, mostly divided into five: Delta, Theta, Alpha, Beta, and Gamma. The decomposition of an EEG timeseries into different bands is illustrated in Fig. 1(a). Each of these bands corresponds to a mental state; for example, the high-frequency Gamma band is associated with the most conscious state of the brain, and the lowest-frequency waves, the Delta waves, correspond to deep sleep.

**Figure 1:**
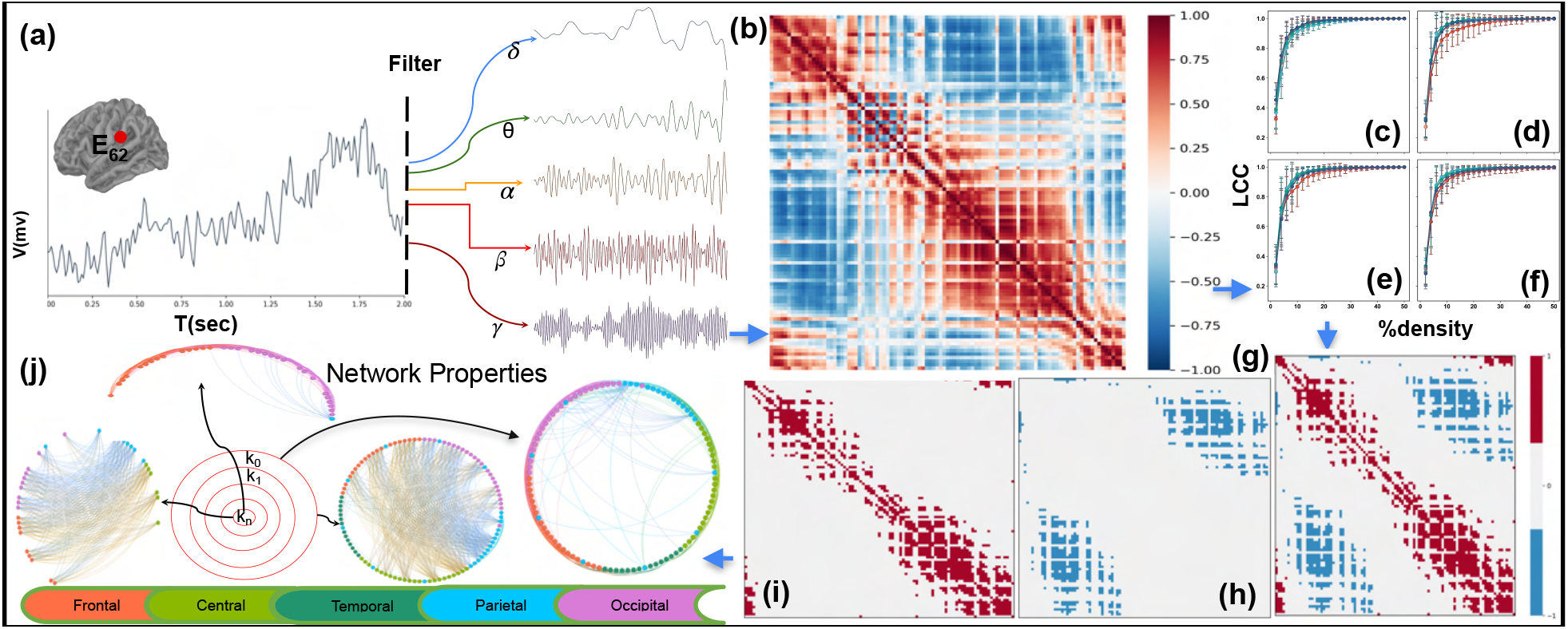
Pipeline for the construction of FCs. The figure presents the pipeline used to study the EEG time series. The preprocessed EEG time series is first filtered to study the different frequency bands separately (a). Once filtered, for each time series we calculate Pearson’s correlation between all pairs of ROIs (b) and then examine the largest connected cluster (LCC) with respect to density (c). The most significant correlations are considered irrespective of the sign. (c), (d), (e), (f) LCC vs Density graphs for resting state, food, image, and word choices, respectively. Based on the LCC and the density plot, we find the density appropriate to proceed with the binarization of the correlation matrix (g). We further divide this matrix into two: one with only negative correlations (h) and the other with only positive correlations (i). Next, we study the network properties of both of these unsigned networks (j).

Furthermore, the EEG-style topographic cortical regions (TCRs) are function-specific: Frontal (F), Central (C), Parietal (P), Temporal (T), and Occipital (O). For example, the Frontal region is responsible for decision-making, and the central region controls motor movements. The Temporal region is involved in memory, face, and object recognition. The Parietal region integrates visual and auditory information, while the Occipital region primarily processes vision[11]. Additionally, all these TMs may also be associated with more complex brain processing, and understanding the human brain and its functioning remains an ongoing challenge in current research.

Brain activity involves both correlations and anti-correlations between different regions (Figs. 2), indicating that both are important. The correlation is associated with the simultaneous involvement in performing a task, as regulated by the underlying physical connectivity between different regions, the negative correlations also play an important role in task-performing states, such as movement control and emotional regulation, and decrease with cognitive load and aging [12–14] and have also been associated with reduced network stability by altering the balance of triads [15]. In this work, we explore in detail the topologies of negative and positive correlations separately to uncover their differences and how their structures change across task states relative to resting states. To achieve that, we construct the functional brain network using Pearson’s correlation, which captures both positive and negative correlations among brain ROIs. We construct the binary functional FCs (details of network generation are in the next section) using a density threshold, accounting for both significant positive and negative correlations. We then separate the positive and negative edges and construct separate functional networks of regions of interest (ROIs) that are positively or negatively correlated, denoted *G*_+_ and *G*_*−*_, respectively. Considering TCRs as the topographical modules (TMs), we find that *G*_+_ functional networks have denser connections within similar TMs than between TMs, consistent with the so far understood modular organization. In contrast, the *G*_*−*_ functional networks have more connections between ROIs across TMs than within TMs, a topology called partially multipartite/anti-modular [16]. The *G*_*−*_ networks have lower density than the *G*_+_ networks, and we find them to be topologically more efficient across all states, highlighting the importance of this topology, as also reported in [16]. Also, we find that the modularity value (see the method section), computed with TMs as modules, is slightly lower than the optimal modularity (without fixing modules). The optimal modularity of the *G*_*−*_ networks also remains positive; however, it is very small, indicating that without fixing the modules as TMs, one cannot find the anti-modular topology. We also study another dataset of eye-open resting-state data to confirm this structure further and obtain similar results.

**Figure 2:**
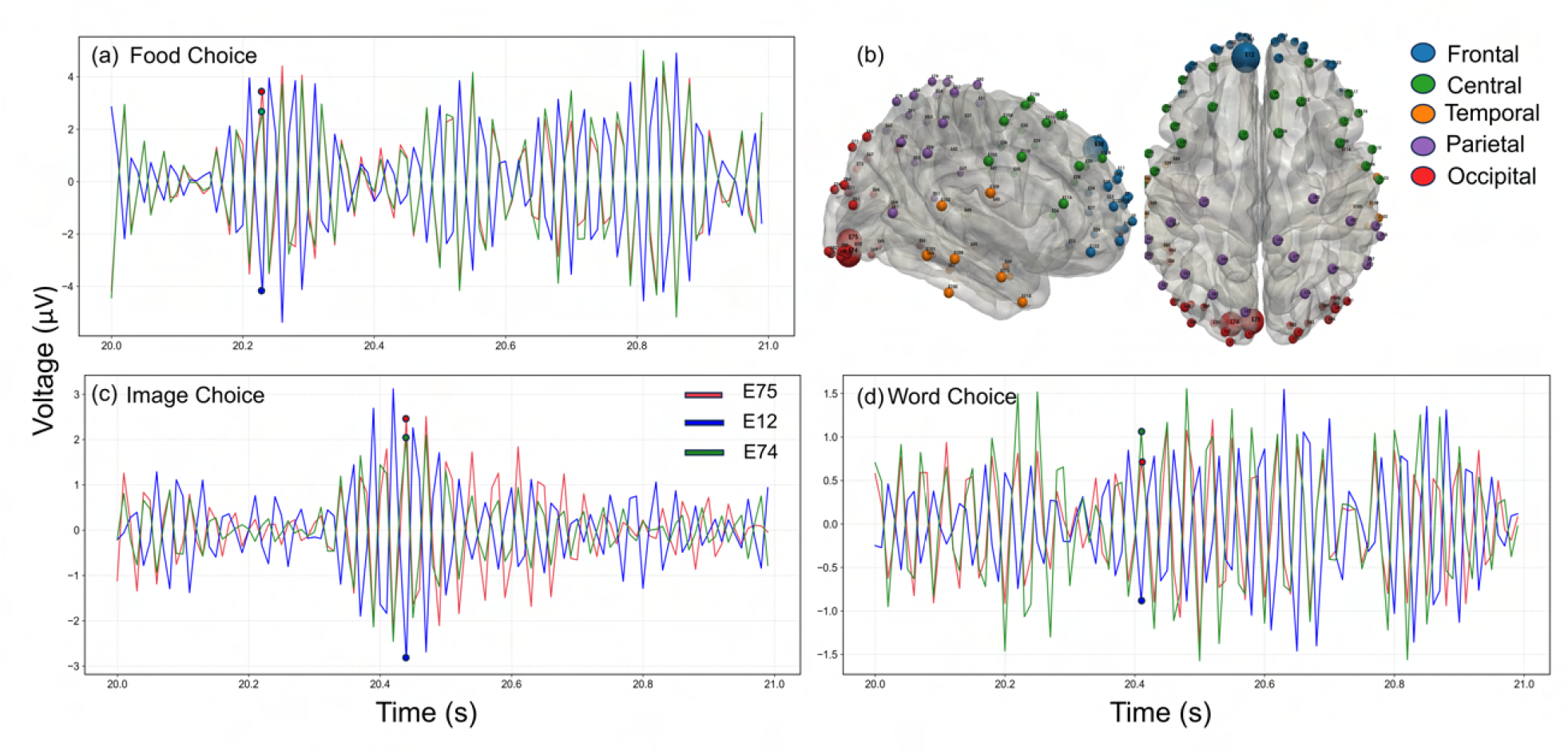
Correlated and anti-correlated ROIs in a brain. Subplots (a-c) present the time series in the gamma band for cognitive task states of a particular subject (Sub-07) for food (a), image (b), and word choices (c) for the E75, E12, and E74 ROIs. (d) shows the position of these ROIs in the brain. Note that the time series fluctuate in their correlation in some instances, but overall the E74, E75 pair is positively correlated, whereas E75, E12 and E74, E12 are negatively correlated.

Further, we perform k-core analysis (see the method section) of all networks, which provides insight into cognitive states and their resilience [2–4]. This analysis provides insights into the robustness of FCs, with stronger core networks associated with better cognitive outcomes [2–4]. Additionally, the core-periphery organization is a stable architectural feature even across neuro-developmental disorders. In the study of Alzheimer’s disease (AD) the left hemisphere k-core is found to be completely missing in AD patients [17]. k-core percolation reveals the hierarchy present at the network level in the structural and the functional networks [18]. We find that *G*_+_ is more hierarchical than *G*_*−*_, as reflected in a greater number of k-shells (see the method section), in both the resting and decision-making states. Also, both *G*_+_ and *G*_*−*_ exhibit greater hierarchy in decision-making states than in the resting state. The mesoscopic structure of the *G*_*−*_ network remains intact at the innermost core level, with the innermost k-core of *G*_*−*_ networks being antimodular; however, it becomes closer to a random network in *G*_+_ networks. The innermost core ROIs also exhibit a distinct spatial distribution. For *G*_*−*_ networks, it is a bimodal spatial distribution for all the bands except the *γ* band in the resting state, for which most of the subjects show a unimodal spatial distribution. However, in the decision-making FCs, it becomes bimodal across all frequency bands. In contrast, the *G*_+_ remain uni-modal for all the frequency bands in both the resting and the decision-making states, reflecting the spatial delocalization of the innermost k-core nodes of the *G*_*−*_ graph. Furthermore, for *G*_+_, the innermost core nodes in the high-frequency *β* and *γ* bands are mostly from the occipital TM in the decision-making. In contrast, the lower-frequency bands, alpha, delta, and theta, mainly derive their innermost core nodes from the central or frontal TM. In the *G*_*−*_ networks, the innermost core shows a significant contribution from ROIs in both parietal and anterior regions. The number of connections within and between the modules of all subjects across all tasks is extensively studied in the high-frequency gamma band in the Fig. S4. In the next section, we elaborate on data acquisition and FC construction.

## 2 Data and Functional connectome generation

The data was collected from [20] https://github.com/andlab-um/MT-EEG-dataset. The data comprise EEG time series from the resting state and the dynamic binary decision processes for semantics and preference choices, collected from 31 subjects performing food-preference and semantic-judgment tasks. It consists of time series from 101 channels, after excluding noise and facial/eye-movement artifacts. The HydroCel 128-channel EEG cap provides 3D coordinates for each node. Following are the details of the experiment performed:

The dataset includes recordings from an experiment consisting of four stages from resting to Food preference, during which subjects are shown 320 food pictures in pairs for selection. Then, an image choice of 160 objects follows, with equal numbers of animate and inanimate objects. The subject is asked to determine whether the shown object is animate or inanimate, and the same objects are then used for Word choice, where images are replaced by Chinese words, with details about the tasks. Data include EEG time series from 129 channels (128-channel cap based on the standard 10*/*20 System with Electrical Geodesics Inc., EGI, Eugene, Oregon). After pre-processing and removing noisy channels, 101 channels are finalized for further analysis. Four subjects, sub-01, sub-05, sub-19, and sub-28, were removed from the final analysis because their performance was poor and data acquisition was noisy. The different channels correspond to different brain regions, commonly referred to as regions of interest (ROI). The brain waves thus obtained comprise different frequencies. A popular division is Delta, Theta, Alpha, Beta, Gamma bands with frequencies (1*−*4 Hz), (4 *−* 8 Hz), (8 *−* 13 Hz), (13 *−* 30 Hz), (30+ Hz) respectively. We then separated the brain waves in the different bands (Fig. 1(a)).

Next, we construct the FC (Fig. 1(b). There are various methods to construct the FC; one method is to compute the linear correlation between the time series of all pairs of ROIs, which is simple yet powerful for extracting meaningful information about functional correlations between different brain regions. The correlation values are interpreted as functional connectivity. The correlations are calculated using Pearson’s correlation (see method section). The correlations calculated thus have both positive and negative values. A positive (negative) correlation between two ROI indicates a positive (negative) linear relationship and functional connectivity between them. Furthermore, because connectivity measures are generally affected by volume conduction, in practice, an appropriate threshold can yield meaningful connectivity. The various methods for thresholding are: absolute thresholding, percolation-based thresholding method, proportional thresholding (density thresholding), minimum spanning tree (MST), and permutation testing [21][22][23]. We chose density thresholding because most network metrics are affected by network density, which can distort decision-making and lead us to miss the actual changes across the states. Implementing density thresholding yields a binary network in which nodes represent different ROIs and edges indicate high functional correlation between them. To find the optimal density threshold value, we begin with a minimum density threshold (1%) based on the strongest correlations. We then calculate the size of the largest connected component (LCC), denoted as (*N*_*max*_). We continue increasing the density until LCC accounted for almost 99% of the ROIs. We observed that the 20% density threshold consistently maintains 99% of ROIs within the LCC across all time series studied. This density threshold ensures the size preservation for the minimum density, keeping in mind that real-world networks are mostly sparse yet connected [24],[25]. We say this matrix is *G*. Further, we divide this matrix into two; one with only positive connections (*G*_+_), and another with just negative connections (*G*_*−*_). We present the entire pipeline of the FC construction in the schematic 1. Moreover, we divide the 101 channels into key segments, viz. Frontal, Central, Temporal, Parietal, and Occipital, based on information provided in [26]. This classification only provides the positioning of the 90 ROIs, and we assign the other 11 ROIs to a segment based on their spatial proximity by calculating the Euclidean distance to their nearest neighbor. This division is confirmed based on the division provided by [27]. The details of the ROIs in different TMs are given in Table S1.

In-addition, to further confirm the anti-modularity and the multipartite nature of anti-correlated (*G*_*−*_) networks, we studied an additional dataset from the eye-open resting-state [28]. We construct the functional network for this dataset as discussed previously, using the same 20% density threshold. In the following, we first study the functional networks *G*_+_ and *G*_*−*_ generated by this method.

## 3 Results

After constructing the FCs from the positive and negative correlations separately, we explore their topology, treating both as unsigned across different frequency bands and states: the eye-open resting state and three binary visual-based decision-making states. In the following, we discuss these results in detail:

### 3.1 The *G*_+_ networks are modular but the *G*_*−*_ networks are anti-modular

We first calculate the modularity values, treating the different topographic cortical regions as distinct modules (see the method section). We refer to this as the fixed modularity (*Q*_fixed_). We compare the empirical networks with the 20-degree-preserving random networks, which serve as the null model (see the method section). Figs.3(b)–(k) present all these analyses. Overall, across the frequency bands (resting or working), the *Q*_fixed_ values for the *G*_+_ networks are positive, but for the *G*_*−*_ networks remain negative. In contrast, the corresponding degree-preserved random networks exhibit values close to zero, indicating a non-random pattern of positive and negative modularity values for the positive and negative correlations between cortical regions, respectively. The negative *Q*_fixed_ values indicate anti-modular connectivity between the five brain segments in the *G*_*−*_ network. In contrast, the positive *Q*_fixed_ values indicate the presence of the modular structure in the *G*_+_ networks. In addition to modularity, we also calculate the Module Connectivity Ratio (*r*) (see the method section) for all FCs and find that it supports the results coming from the modularity values (Figs. S1, and S2). Our results indicate that most subjects in high-frequency bands exhibit an additional layer of complexity, characterized by intra-TM negative connections in both resting and task states. Fig. S4 further illustrates this point, emphasizing that the within-module negative connections are more in the resting states and within all TMs, but in the task states, these connections are very few or absent in most subjects within the temporal, occipital, and parietal regions. This indicates that the task states correspond to the strongest positive correlations in the temporal, occipital, and parietal regions, reflecting greater functional specialization in these regions.

In contrast, the low-frequency bands in the resting state do not exhibit the same level of complexity due to the absence of the within TM negative correlation and this results in the existence of the multipartite (rather than partial) structures in most of the subject’s *G*_*−*_ networks.

Moreover, the *Q*_fixed_ for the overall FC (unsigned *G*) takes only positive values, and the corresponding degree-preserved random networks exhibit very small, near-zero values, indicating stronger positive correlations between ROIs within the cortical regions than the negative correlations between the ROIs in different TMs. In terms of network property, *G*_+_ networks possess higher density than *G*_*−*_ networks (further demonstrated in fig.4). Note that the *Q*_fixed_ values for the *G* networks are much lower than those for *G*_+_, showing the negative correlated inter-TMs connections are diluting the modular structure of the positive correlations when they are considered as a single (shown in gray box plots in Figs. 3( b, d, f, h, and j)).

**Figure 3:**
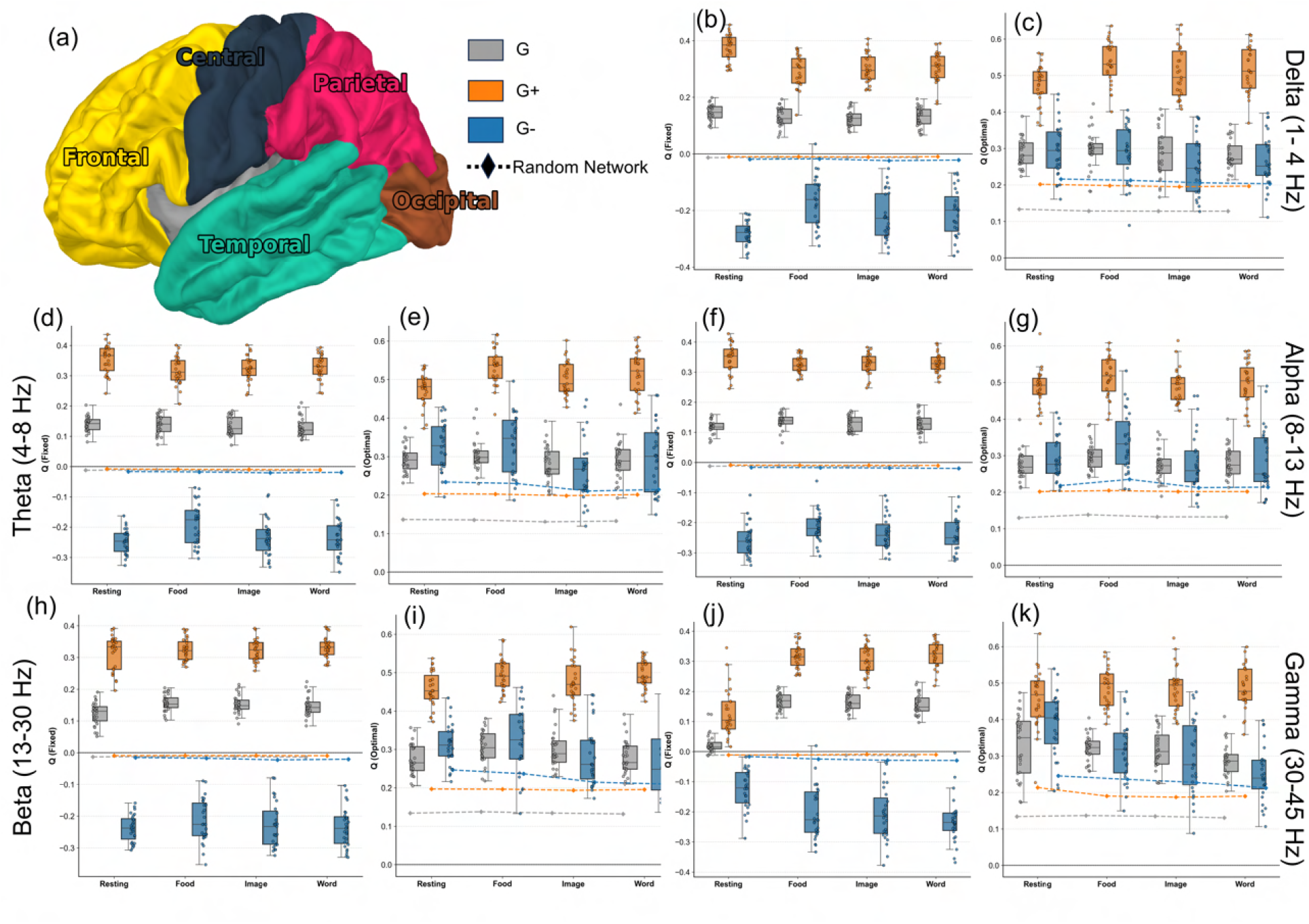
Modularity and Anti-modularity in FCs across different states and frequency bands. Subplot (a) displays the EEG-specific TMs (adopted from [19]). Subplots (b, d, f, h, j) plot of fixed modularity (*Q*_fixed_) for all five frequency bands across three network types, considering TMs as modules. The optimal modularity (*Q*_optimal_) for all networks is shown in the subplots (c, e, g, i, k). All the subplots (b-k) show the comparison of modularity values with those of corresponding degree-preserving random networks (dotted lines). The positive (negative) values of the *Q*_fixed_ indicate the existence of the modular (anti-modular) topology in these networks with five TMs as modules; the value of (*Q*_fixed_) without sign shows the strength with which these properties are preset in a network.

Furthermore, we study an additional eye-open resting-state data set and find that, when considering TMs as modules, *G*_+_ networks exhibit a modular structure (Fig.S3). In contrast, the *G*_*−*_ networks exhibit an anti-modular structure (Fig.S3), further confirming the modular and anti-modular structures of the *G*_+_ and *G*_*−*_ networks, respectively.

We also study the optimized modularity (*Q*_optimal_) using Newman’s method [29] (see the method section) to examine how the functional topographic modules (TMs) differ from the functional modules obtained via modularity optimization of binarized correlation networks. *Q*_optimal_ remains positive for all, *G*_*−*_, *G*_+_, and *G*, showing correlation-based optimal modularity in the studied FCs (Figs.3 (c), (e), (g), (i), (k)). *Q*_optimal_ remains higher for the empirical networks than the corresponding degree-preserved random networks. Besides, the *Q*_optimal_ always takes higher values than the *Q*_fixed_ for all the studied networks. Owing to this, though the brain segments Frontal (F), Central (C), Temporal (T), Parietal (P), and Occipital (O) collectively perform specific cognitive, sensory, and motor functions, in the resting and decision-making brain, they do not form optimal modules. Different ROIs within these segments can exhibit greater connectivity than those within the same segment.

Next, comparing FCs across the frequency band, we find visible changes in the resting and decision-making brains. The *G*_+_ networks show a decrease in *Q*_fixed_ values in the task states compared with the resting state in the low-frequency delta, theta, and alpha bands. In the beta band, the *Q*_fixed_ value for the resting state and the decision-making states becomes almost similar, whereas in the high-frequency gamma band, the *Q*_fixed_ value for the task states is significantly higher than the resting state (Fig. 3(i)). Indicating that topographical modules exhibit stronger correlations within themselves in the task state compared to the resting state in the high frequency bands. This stems from enhanced intra-TM connections across all TMs, with the effect more pronounced in the occipital region (Fig. S4). In the low-frequency band, the resting-state has more functionally segregated TMs than the task-state, suggesting that a calm brain mostly operates in this frequency regime. Similarly, |*Q*_fixed_| for the *G*_*−*_ networks is significantly higher for the task state than the resting state in the high-frequency gamma band. In contrast, in the alpha and beta bands, it remains slightly lower, and in the low-frequency bands, it is significantly lower than the resting-state value. Moreover, *Q*_optimal_ in *G*_+_ also exhibits higher values in the task state than the resting state in the gamma band and beta band, and similar or higher values in the lower frequency bands. The *G*_*−*_ does not exhibit a similar trend: it shows lower *Q*_optimal_ values in the task state than in the resting state in the high-frequency bands (Figs. 3(k)).

#### 3.1.1 Global integration, segregation, and Positive to negative Edge ratio

To further explore the impact of the coexistence of modularity and anti-modularity within topological modules on the overall performance of the different networks, we examine the global efficiency and average clustering coefficient (see the method section) across all empirical networks. To compare them with the null models, we calculate the efficiency and the global clustering for their corresponding degree-preserved random networks. We plot the ratio of the efficiency and clustering of empirical to the random in Fig.4 (a) and Fig.4 (b), respectively. We find that the global efficiency is considerably lower in *G*_+_ networks than in *G*_*−*_ networks, and that the latter have a higher global efficiency, closer to that of the corresponding degree-preserved random networks. Moreover, the anti-modular *G*_*−*_ networks have almost negligible clustering as compared to the modular *G*_+_ networks (Figs. 4(b)). The *G* network, which combines *G*_+_ and *G*_*−*_, exhibits both higher efficiency and clustering, with efficiency exceeding that of the modular *G*_+_ networks and clustering exceeding that of the *G*_*−*_ networks. The clustering of all the networks *G*+, *G*_*−*_, and *G* differs significantly from the resting state to the task state in the higher frequency bands, and the resting state is less clustered than the task state.

**Figure 4:**
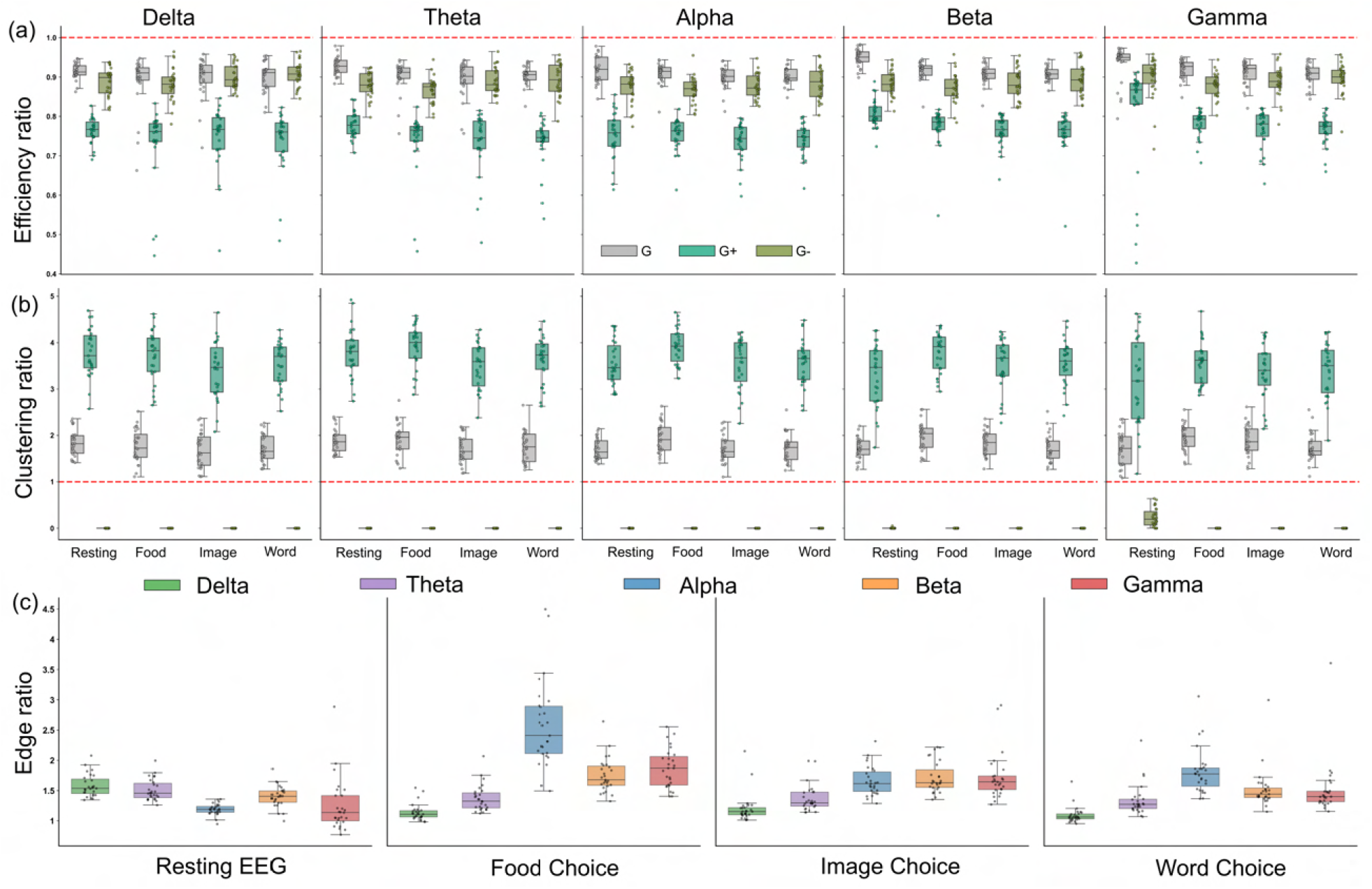
Network properties of the FCs across different tasks and the frequency bands. (a) The box plots of ratios of Global Efficiency(*E*) values for empirical and random networks for all subjects in all three types of FCs across different bands. (b) plots ratios of Average global Clustering Coefficient (*C*) values for empirical and random networks for all subjects. (c) plots the ratio of positively (*N* ^+^) to negatively (*N* ^*−*^) correlated ROIs across all tasks and frequency bands; the open circles are statistical outliers. The red-dashed line at *Y* = 1.0 represents the theoretical baseline for the FCs having a random structure.

Furthermore, the significant negative correlations are lesser than the positive correlations evident from the positive (*N* ^+^) to negative (*N* ^*−*^) edge ratio plots in Fig. 4(c). So, *G*_+_ is always denser than *G*_*−*_. Network metrics, such as efficiency and clustering, are affected by density: a denser random network is more efficient than a sparser one. However, despite *G*_+_ being significantly denser than *G*_*−*_ networks, the efficiency of the *G*_*−*_ networks is much higher than that of the *G*_+_ networks. Showing that the anti-modular organization imparts better efficiency to the network than the modular organization. However, higher clustering is also associated with higher modularity, which imparts segregation and, hence, functional specialization. The observation of anti-modular negative correlations suggests that the brain has to organize to exhibit this anti-modular nature in the negative correlations between topographical modules, which are functionally specialized anatomical modules. Moreover, we observe an increasing trend in median values across low-to high-frequency bands in task states. The high-frequency bands (e.g., gamma) exhibited a larger IQR and more distinct boundaries than low-frequency bands. The alpha band also showed a slightly higher interquartile range (IQR) and distinct boundaries, indicating subject-level variability in decision-making during food selection. In contrast, lower-frequency bands showed no such differences across states.

## 4 k-Core-analysis

We perform *k*-core analysis (see the methods section for details) on *G*_+_ and *G*_*−*_ networks separately. K-core percolation reveals the hierarchical organization and robustness of networks. We separately calculate the fraction of ROIs in each *K*-core of the *G*_+_ and *G*_*−*_ networks. We then average the fraction of ROIs for each K-core across subjects, tasks, and frequency bands. Fig.5 plots all this analysis. We find that *G*_+_ and *G*_*−*_ behave differently to the k-core pruning; The *G*_+_ graphs display a more gradual decay with a longer tail (Figs. 5(a)). Whereas, the *G*^*−*^ graphs exhibit a steeper and faster decay with a short tail (Figs. 5(b)). Moreover, there is a difference in the pruning of resting-state and task-state graphs across frequency bands. The alpha band of the resting-state in the *G*_+_ network has a slightly shallower decay curve, whereas the remaining bands decay significantly faster. Across task conditions, the Image Choice and Word Choice tasks show similar decay curves and patterns across all frequency bands, especially in *G*_+_ networks. Further to quantify the k-core percolation robustness, we calculate the area under the k-core decay graphs (AUC), as shown in Fig. 5(c). In the *G*_+_ network, the mean AUC during task states consistently equals or exceeds that of the resting state, with a significantly higher value in the Gamma band. In the *G*_*−*_ networks, the median AUC in the resting state is slightly higher in the low-frequency delta and theta bands and is lower than or equal to that in the task states in the higher-frequency bands. Furthermore, the innermost K-core numbers are also studied in Fig.5(d), and the trend remains the same: task states have higher core numbers, indicating that many ROIs survive the pruning process and attain higher core numbers than in the resting state. Meanwhile, except for the dip from resting state to food choice in the alpha band, the *G*_*−*_ network shows a similar trend, with lower values. Collectively, it indicates how robust the respective networks are during task performance and how cognitive load drives them to engage deeper core networks. The lower median AUC in the gamma band for food choice than the resting state, but the higher *k*_max_ value indicates that, in the k-core percolation state related to food choice, it is less robust in terms of k-shell occupancy but is more hierarchical in constituting the ROIs surviving for deeper k-values.

**Figure 5:**
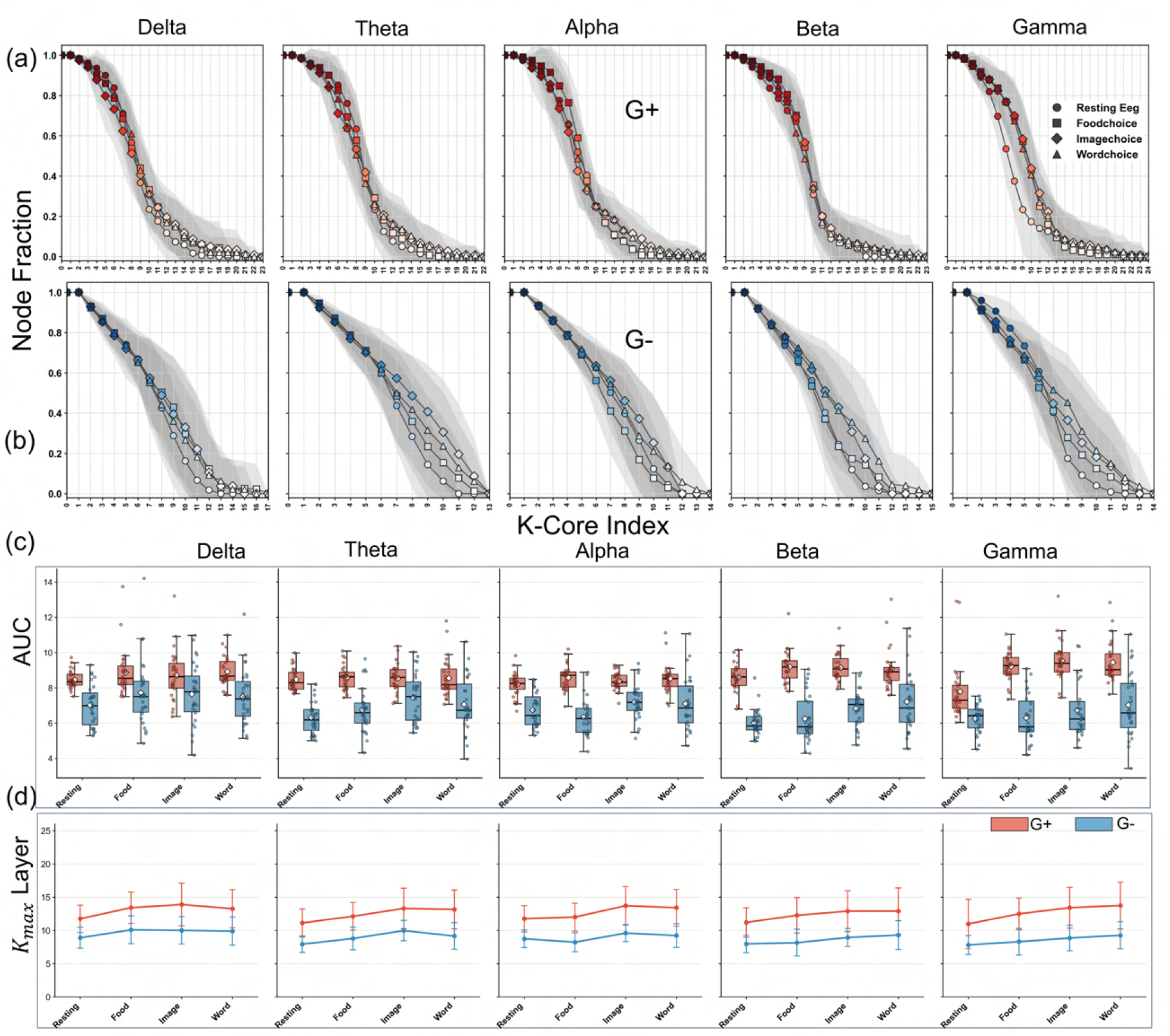
K-core analysis of FCs across different states and frequency bands. Subfigures (a) and (b) display the k-core decay (also called k-core percolation) graphs of the *G*_+_ and *G*_*−*_ networks, respectively, across various frequency bands. These plots show the mean fraction of ROIs with core number *≥ K* across subjects, with the grey-shaded regions denoting *±*1 standard deviations from the mean. The change of color from thick red/blue to white also indicates the fraction of ROIs in different subjects. *G*_+_ networks show more gradual decay with longer tails and high core layer values, whereas *G*_*−*_ networks are steep with minimal core penetration. Subplot (c) plots the area under the curve of K-core percolation graphs. Finally, subplot (d) plots the maximum core index (*K*_max_) with error bar across subjects.

The image and word choices yield closer k-core pruning graphs across all frequency bands. We can interpret this similarity in light of the experimental design: the Image Choice task requires subjects to classify objects as animate or inanimate, followed by the Word Choice task, in which the same objects are presented in textual form and reclassified. Plausibly, because subjects have already been exposed to visual representations of the objects, the neural processing required for the word-based classification is likely to engage overlapping cognitive mechanisms[30]. Moreover, overall, this analysis hints that the decision-making states are more robust than the resting state. The lower robustness of the resting state compared to task states also suggests that focused task states are more robust.

### 4.1 Innermost core ROI and their connectivity

In the k-core decomposition, the innermost core members form the most well-connected base of the Hierarchy, reflecting their importance in overall interactions between different ROIs and in the processing occurring in a particular brain state. In the following, we look into their composition and the spatial localization, and discuss these results in detail:

#### 4.1.1 Spatial localization/Delocalization of the innermost core ROI

The time series plotted in Fig.2 show that the ROIs from the same TM (occipital in the figure) exhibit in-phase spiking, whereas pairs from different TMs (occipital and frontal in this figure) exhibit out-of-phase voltages, hinting at negative correlations between the distant regions, which we confirm by finding the Pearson’s correlation between them as mentioned in previous sections. The time series of these ROIs are also plotted, considering them part of the innermost core in most subjects and across the high-frequency beta and gamma bands. In this section, we explore whether the innermost core exhibits such nonlocalization and whether the innermost core of positive correlations exhibits a similar behavior. To understand the localization/delocalization of innermost core *k*-core ROIs, we first visualize their pairwise Euclidean distances.

We plot the pairwise distances between the core ROIs of one brain (subject 7 in the data studied) in Fig.6(a,b). We observe a bimodal distribution in the innermost core of the *G*_*−*_ graphs across all bands and tasks. Whereas, for the *G*_+_ graphs, although the variance is high, the distribution appears unimodal. To rigorously quantify this visually observed spatial divergence, we used Sarle’s Bimodality Coefficient (BC) [31] (see method section). We calculate BC utilizing the sample skewness and sample kurtosis of the distance distribution. Following [31], a threshold of *BC >* 0.555 statistically confirms the bimodal or multi-modal distribution. We plot BC for all 27 studied subjects in Fig. 6(c) and (d) for *G*_+_ and *G*_*−*_ graphs, respectively. The BC values for the *G*_+_ graphs remain below 0.55 across all subjects, frequency bands, and states; however, the *G*_*−*_ graphs exhibit different behavior. For *G*_*−*_ graphs, the median of all the states and frequency bands remains greater than 0.555 except for the resting state Gamma band, for which some of the subjects show values greater than 0.555, but mostly show values less than 0.555. This indicates that in *G*_*−*_ networks, delocalized spatial hubs form, suggesting that the innermost core ROIs are contributed by brain segments that are not close to each other. In contrast, a unimodal distribution in *G*_+_ networks can be explained by ROIs arising from spatially closer brain segmentation, so that the respective pairwise distances are considerably small. Moreover, we calculate the mean pairwise distances for all the brains and find that the mean for the *G*_+_ networks has more spread than the *G*_*−*_ networks (Fig. 6(e) and (f)), showing that the overall average distance between the innermost core neurons of the different subjects is more similar than the *G*_+_ network, indicating that the subjects are more similar in terms of the *G*_*−*_ networks. In contrast, based on the *G*_+_, their subjects are more different.

**Figure 6:**
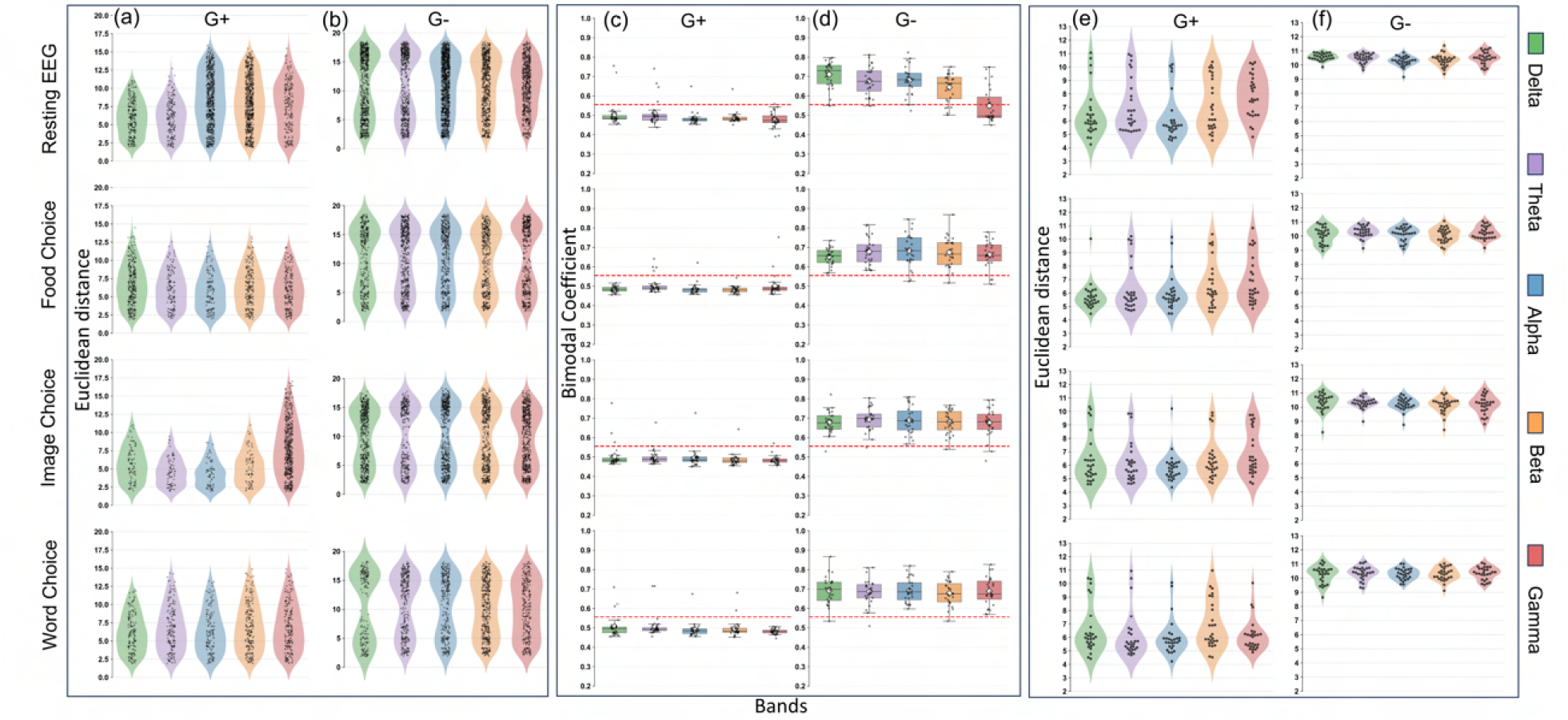
Spatial localization and delocalization of the innermost core of *G*_+_ and *G*_*−*_ networks.. Subplots (a) and (b) demonstrate the innermost core ROIs’ pairwise Euclidean distances for one of the subjects, Sub-07 and (c),(d) plots of Bimodal Coefficient values for 27 subjects across all tasks and bands. The diamonds in the box plot display the mean values. The red line corresponds to the Bimodality coefficient value 0.555, a value above which demonstrates the existence of the bimodality in a given distribution. Finally, the Violin plots in (e) and (f) display the mean pairwise Euclidean distances for all ROIs of the brain of all the subjects.

#### 4.1.2 Topology of the innermost core ROIs

We further explore the topology of the innermost k-core ROIs. First, we calculate the topographical module connection probability (see methods section) for inter- and intra-topographical module connections and find the ratio of these (*r*). *r >* 1 indicates more connections between the topographical modules than within them, i.e., an anti-modular topology. The *r* values less than 1 indicate that the connection probability between modules is greater than the probability between modules, indicating the existence of a modular topology. The *r ≈* 1 shows the existence of the random topology. Fig.7(a) plots these values for all the subjects for all the states and the frequency bands. We find that for the *G*_+_ networks, the *r* values for most of the subjects take values close to 1, whereas for *G*_*−*_ networks *r* takes values much greater than 1, indicating that the innermost cores of the *G*_+_ networks are randomly connected. In contrast, those of the *G*_*−*_ networks form the anti-modular networks. Indicating that for the *G*_*−*_ networks, even though the *k*_*max*_ core makes up a relatively small fraction of the whole network, there is a resemblance between the brain’s innermost core and its global network structure (Fig. S1). Next, to explore the association of the innermost core ROIs of the *G*_+_ and *G*_*−*_ networks to different TMs. First, we plot these networks for a brain (subject 7 of the data set, Fig.7(b), (c)). It shows that the inner-core ROIs of *G*_+_ are located in either posterior or anterior brain regions. In contrast, the innermost core ROIs of *G*_*−*_ are distributed across both anterior and posterior brain regions. Next, we determine the fraction of ROIs in the innermost core across all subjects and TMs and again, we find that the *G*_*−*_ and *G*_+_ networks exhibit distinct characteristics. The *G*_+_ networks exhibit a higher contribution of the anterior regions than posterior in the alpha band (Fig. 8(a)), which remains true for the other lower frequency bands too (Fig. S5); however, in the high frequency gamma band, it shifts to the posterior regions. Whereas the *G*_*−*_ networks, exhibit a balanced or relatively greater contribution from the posterior regions than from the anterior regions in all frequency bands, both in the rest and in the task states. Moreover, the balance between the contribution from the frontal to occipital in *G*_*−*_ networks in the innermost core is more prominent in the resting state alpha band than the task states; however, in the gamma band, it is more prominent in the food and image choice tasks than the resting state, and in all the states and bands there is less variation across the subjects (Fig. 8(b))(for remaining band details see S6). Whereas in *G*_+_ networks, there is huge heterogeneity between this ratio across the subjects in the alpha band, which narrows down in the gamma band (Fig. 8). The lower heterogeneity of the subjects in the gamma band reflects a similar processing mechanism across the different brains. Among all the task states in the gamma band, the image-choice task shows greater heterogeneity across subjects.

**Figure 7:**
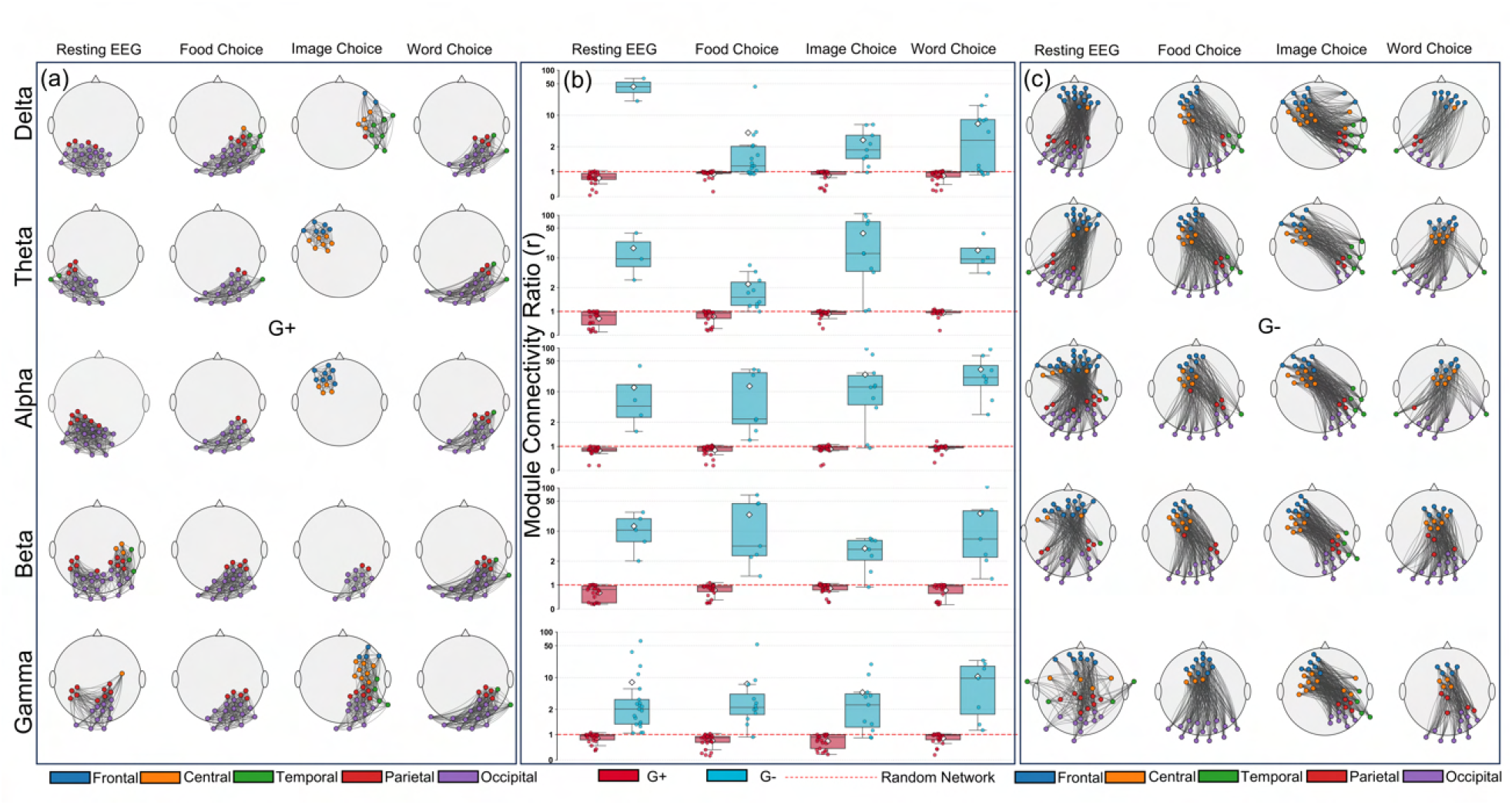
Topology of the innermost core of *G*_+_ and *G*_*−*_ networks. The subplots (a-c) demonstrate the topology of the innermost core ROIs of the *G*_+_ and *G*_*−*_ networks. (b) plots Module connectivity Probability ratios (r) of all 27 subjects. The *r* value for the innermost core of some subjects takes very high values; therefore, a logarithmic scale is used on the Y-axis from *r* = 2 to *r* = 100. The plots (a) and (c) exhibit the topology of the innermost core of *G*_+_ and *G*_*−*_ networks, respectively, for one of the subjects (Sub-07) across all the bands and states.

**Figure 8:**
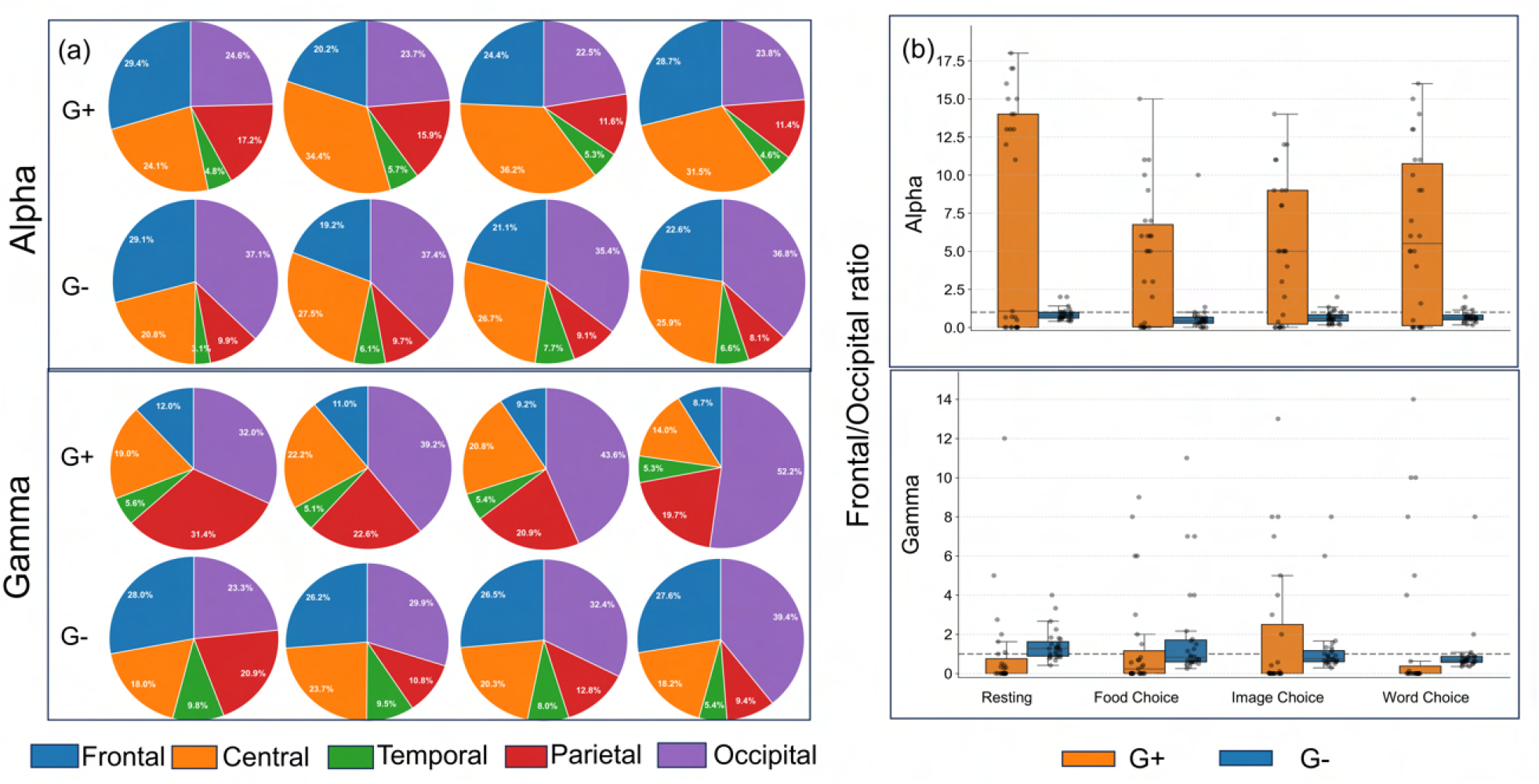
Innermost core ROIs in different TMs. Subplots (a, b) display the distribution of innermost core ROIs in the 5 TMs in alpha and gamma bands, respectively. In generating this figure, the innermost core ROI for each subject is counted, considering an ROI as many times as it appears as an innermost core ROI; similarly, the total number of ROIs is counted across subjects. Subplots (c, d) demonstrate the fraction of the frequency of frontal to the occipal innermost ROIs.

#### 4.1.3 Spread of innermost ROI across subjects and relative topological shift from Rest to task

Further, to explore the contribution of the different TMs in the innermost core of the different subjects in the two graphs, we find the union of innermost core ROIs of all subjects, divide them to different topographical modules and and plot the fraction of times they appear as in inner most core of different subjects (shown as box plots in Fig. 9).

**Figure 9:**
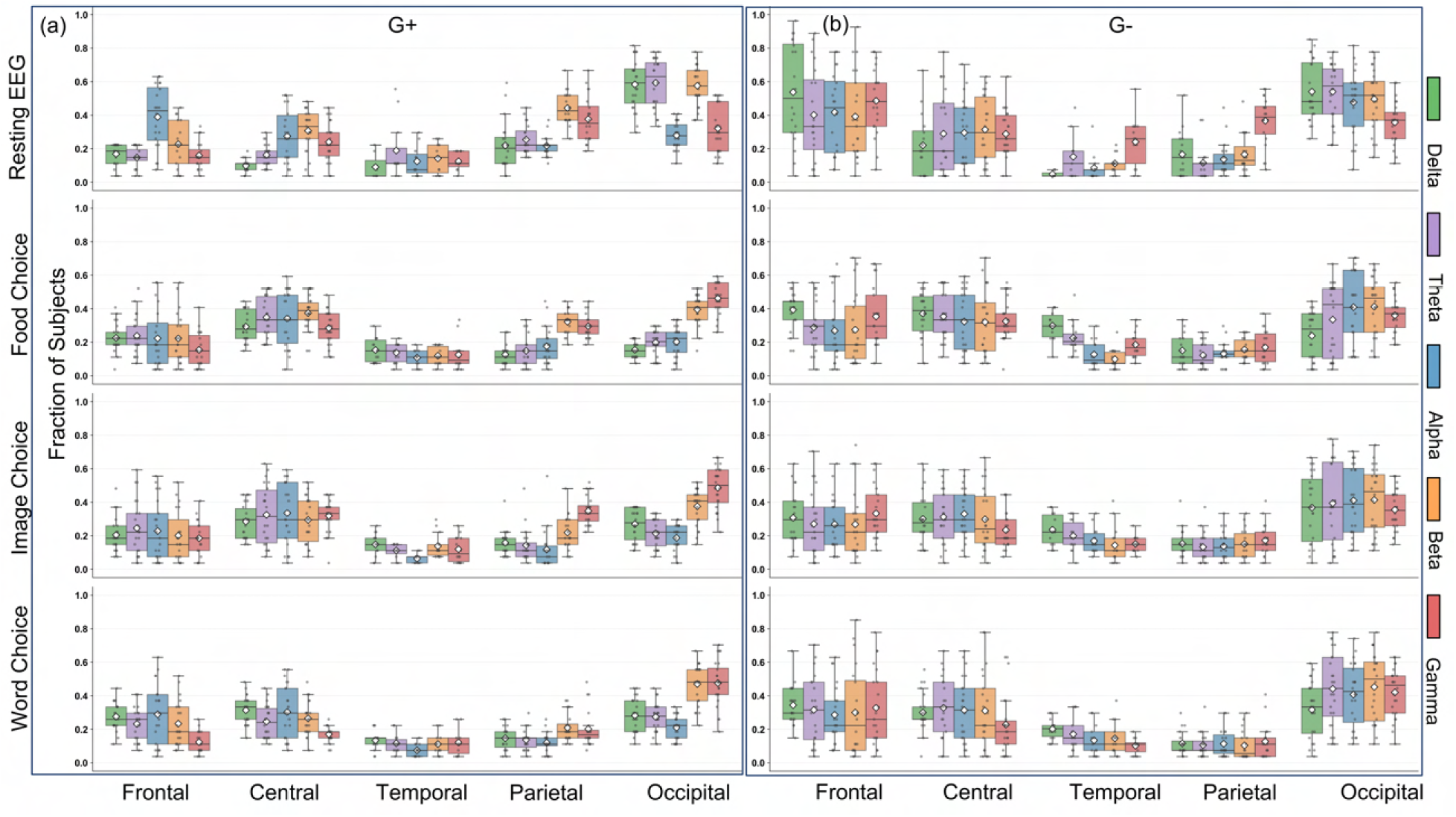
Variation of the innermost core ROIs across the subjects. This figure shows the fraction of subjects with a common innermost ROI. We plot this fraction for the different ROIs and TM groups separately. White diamonds denote the mean, horizontal lines denote the median, and individual grey scatter points represent specific ROIs within the segment.

When it comes to ROI consistency across subjects in terms of their occurrence as the innermost core ROI, it may differ across frequency bands and states. In *G*_+_, the ROIs from the frontal lobe in the alpha band show a drastic decrease in their consistency across subjects. The occipital ROIs exhibit a similar trend in the alpha, theta, and beta bands. The consistency of the parietal ROIs across subjects decreases across all bands except the alpha band, which shows greater consistency in task states. In the *G*_*−*_ networks, the frontal ROIs show decreased consistency across subjects in all bands. In contrast, the occipital ROI shows a decline in consistency in all the bands and task states from the resting state, except in the word choice task, in which one occipital ROI shows a slight enhancement in the fraction of subjects having in the innermost core. Moreover, temporal lobe ROIs show greater consistency across subjects in the delta band during task states. Overall, the resting state shows the highest fraction of subjects with similar ROIs in both the *G*_+_ and *G*_*−*_ networks, with this phenomenon being prominent in the *G*_*−*_ networks.

Moreover, the inter-quartile range (IQR) of the box plots in Fig. 9 reveals the topologically heterogeneous nature of ROIs across subjects. In the resting *G*_+_ network, the anterior alpha band is wide, but it narrows in posterior regions. The task states in the anterior regions have shown greater spread in both the theta and alpha bands. Conversely, in the high-frequency gamma band during task performance, the lower IQR than in the resting state reflects less heterogeneity in both posterior and anterior regions. The *G*_*−*_ networks also exhibit a very high IQR in the resting state compared to the task states across all bands, in both the anterior and posterior regions. During tasks, there is compression and a median drop, mainly in the low- to mid-range frequencies in the frontal TM. Whereas the posterior regions still show wider IQRs across all frequency bands of task states, which could be due to subject-specific differences in the occipital ROIs’ filtering of background noise and to differences in attention span across subjects.

### 4.2 Comparison between the tasks

The mesoscopic study revealed interactions between cortical regions and their presence in the innermost cores during cognitive tasks in the correlated and anti-correlated networks. We see substantial changes in topology and robustness in the task state from the resting state; however, this does not give a clear difference between the task states of the vision-based binary choices. Therefore, to further achieve this, we delve deeper into the core analysis and examine the intersection ROIs from the top 10 innermost core ROIs across the beta and gamma bands. These are plotted on a 3D brain image for each cognitive task for both the *G*_+_ and *G*_*−*_ networks in Fig. 10(a,b). The *G*_+_ network shows intersecting ROIs from the occipital region, whereas in the In the *G*_*−*_ network, frontal and occipital regions contributed to the intersection of the top 10 innermost core ROIs in the beta and gamma bands, further demonstrating the multipartite nature of *G*_*−*_ networks. The occipital regions are present across most subjects and across the beta and gamma bands in the correlation network, indicating that these ROIs play an important role in processing visual information and in subsequent decision-making. The ROIs *E*_66_, *E*_70_ and *E*_74_ (Fig.10(a))obtained for the Image processing has overlap with both the word choice and the food choice, showing that these ROIs has a special role in processing the optical information within the occipital lobe and between the other TMs.

**Figure 10:**
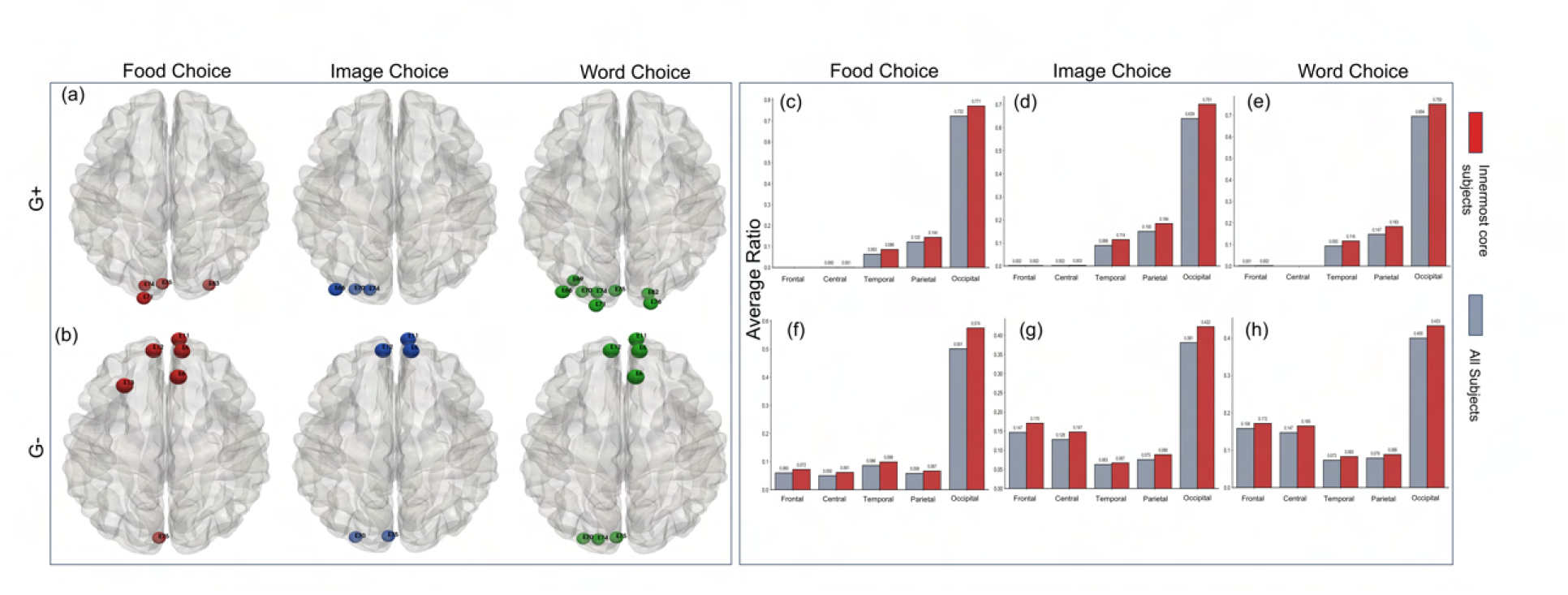
Innermost consistent ROIs across most subjects and beta and gamma bands. The subplots (a) and (b) plot the dorsal view of an intersection of ROIs from the top 10 innermost ROIs found in all subjects across the beta and gamma bands in correlated and anti-correlated networks, respectively. Next, the subplots (c)–(h) illustrate the fraction of affiliations to different TMs for the first neighbors of the ROIs shown in (a) and (b), respectively. Since not every subject has these ROIs in their innermost core, separate ratios are also presented, including all subjects regardless of whether the ROI’s present in the innermost core.

Further, to better understand the functionality of these ROIs, we examine their first neighbors within the respective *G*_+_ and *G*_*−*_ networks. The intersection ROI may not be present in every subject, so we plot the neighbors of the ROIs in subjects with them in the innermost core and those without them in the innermost core separately. The difference is not so significant as shown in Fig.10(c,d,e,f,g,h). However, we find a significant difference between the *G*_+_ and *G*_*−*_ networks; the ROIs of the *G*_+_ network are mostly connected to the occipital region, which is not surprising, given that we already observed the *G*_+_ network to have a modular structure. These ROIs have next-neighbors mostly from occipital regions, then from temporal and parietal regions, but with almost negligible connections to the anterior frontal and central ROIs. Showing not only that the innermost ROIs are localized, but also that the connections of the consistent ones are also localized and do not cover the entire brain. Again, there is not much difference between the connectivity of the innermost ROIs of the different task states, other than the fact that the ROIs for the food choice have fewer connections with the temporal and parietal regions than the image choice and word choice, indicating that image choice and word choice require more correlations between the occipital and temporal and occipital to parietal regions. The temporal regions are found to be associated with visual memory storage, and they are more closely linked to occipital regions; tasks associated with image and word choice are more complex than those associated with food choice. The consistent ROIs in *G*_*−*_ networks are distributed across both frontal and occipital regions and, consistent with the global multipartite nature, exhibit significant connectivity with posterior regions. However, this connectivity is stronger in the anterior regions. Moreover, these show significant differences between the food choice and the other two task states. In food choice, only one occipital region is common innermost ROI in beta and the gamma bands of a significant number of the subjects, which becomes two in the image choice and three in the word choice, showing a more significant contribution of the negative correlations between the frontal and occipital regions in the word choice than in the image and food choice. Moreover, the *E*_75_ channel from the occipital region is common across all tasks. Similarly *E*_12_, *E*_11_ and *E*_5_ are also common across the different tasks. These ROIs play an important role in maintaining both correlations and anti-correlations during task execution based on visual inputs. Moreover, consistent ROIs in the image choice and word choice have better connectivity with the frontal regions, which can be understood from the more ROIs being in occipital for these; however, they have more connections with other TMs too than the food choice task, further explaining the complexity of the image choice and word choice tasks than the food choice tasks.

## 5 Discussions and conclusion

The complexity of neural connectivity provides an infinite number of efficient cognitive states that correspond to both correlation/coherence and anti-correlation/incoherence between different brain regions. EEG time series analysis and network theory are strong pillars of computational neurobiology for extracting information about the patterns of these correlations. Here, we select the publicly available data set comprising time series from the eye-open resting state and three binary decision-making states to explore the topology of the positive and negative correlations, and changes in task states relative to the resting state. Considering five TMs as modules, we find that the *G*_+_ networks comprising positively correlated ROIs exhibit a modular network topology. In contrast, the *G*_*−*_ networks comprising negatively-correlated ROIs exhibit an anti-modular (partially multipartite) topology. To further confirm the partial multipartite (anti-modular) structure of the anti-correlated networks, we also study another publicly available dataset from the eye-open resting state and find the same structure. We also study optimal modularity and find that it is higher than the fixed modularity across all frequency bands in both resting and task states, with task states gaining higher modularity than rest states. The modular organization for the structural connectome is needed for cost-efficiency while maintaining overall connectivity and integration [18, 32, 33]. A correspondence between the functional and structural connectome supports the modular organization of the functional connectomes [34]. However, the functional connectomes also display anti-modularity, indicating that even though the structure corresponds to the so far understood modular organization of the brain, different functional brains may coexist with both modular and anti-modular topologies. The coexistence of anti-modulatory structures in the FCs also reflects the brain’s underlying mechanism for maintaining the functional specialization of the TMs.

Moreover, the topographical modules exhibit stronger positive correlations within themselves in the task state, reflected by enhanced modularity, than in the resting state, particularly in the high-frequency bands. The enhanced modularity also corresponds to almost no negative correlations within parietal, temporal, and occipital regions. Indicating the task states required more in-phase coordination within these TMs. In contrast, in the low-frequency bands, the resting state has more functionally segregated topographical modules than the task state, suggesting that, in the absence of a focused task assignment, the eye-open resting state primarily operates in the low-frequency regime.

Further, *K*-core pruning distinguishes the *G*_+_ and *G*_*−*_ networks, with the former having more *K*-shells than the latter, implying a more hierarchical structure for the former than for the latter and indicating that the positive correlations form a larger *K*-value backbone and a *K*-core structure than *G*_*−*_ networks. Besides, the task states have more *K*-shells and a greater area under the *K*-core percolation graph than the resting states in both graphs, indicating that, in the task state, the brain shifts from relatively fragile to robust by dynamically wiring itself into a deeply hierarchical core-periphery structure designed to handle heavy cognitive lifting. This also suggests that the task state requires more robust, hierarchical functional connectivity for precise decision-making than the resting state, where no specific output is expected.

Furthermore, the innermost core of the *G*_+_ networks is randomly connected and localized. In contrast, in the *G*_*−*_ networks, the innermost cores are anti-modular and delocalized, involving ROIs from anterior and posterior brain regions. The localized topological backbone of the *G*_+_ networks serves as the processing center for positive correlations arising from spatially close brain regions, particularly in posterior regions, during the task state. Whereas the backbone of the negatively correlated network arises from both anterior and posterior brain regions, exhibiting a strong requirement for negative correlations for overall brain functioning across all states. The observation that frontal regions are strongly negatively correlated with occipital regions indicates that the anticorrelations mostly drive the long-range connections responsible for global integration. Besides, the *G*_*−*_ networks exhibit relatively less variability across subjects than the *G*_+_ networks, indicating that the topology of negative correlations is more universal than that of positive correlations across subjects. Additionally, finding that the posterior occipital region is common across all task states indicates that these ROIs are the topological center for both positive and negative correlations, playing an important role in receiving, processing, and communicating visual inputs with other brain regions, as further processing and decision-making require some of these ROIs to play a central role. The strong anticorrelation between these and the frontal region reveals an important brain mechanism that inhibits unnecessary visual inputs and supports greater functional specialization of the different TMs. Moreover, the negative correlation, which can also be interpreted as a lagged correlation between the innermost occipital regions and the frontal regions, indicates that their functioning is interdependent, causing a lag in their firing.

Overall, our analysis of FCs across various tasks and the rest state highlights the mechanisms the brain employs in each process, finds them to be frequency-band-specific, and suggests they may also be subject-specific. Altogether, it points out that while positive correlation within TMs remains the mechanism underlying functional specificity, negative correlation between them is also one of the brain’s mechanisms for specialization and overall functioning, and the functional brain coexists with both modularity and anti-modularity. Moreover, the observation that in task states the most common innermost ROIs of the network with negative correlations across subjects and high-frequency bands are in the frontal and occipital regions, whereas for the networks with the positive correlations it is the occipital region, indicates the central role of the occipital region in possessing and integrating overall input, primarily by exhibiting significant lagged coordination with the frontal region in the tasks based on visual inputs. Our work provides a basis for understanding the topology of negative correlations in FCs and how it differs from that of positive correlations as states shift from resting to task. It ignites further curiosity about the functional brain’s topology of negative correlations in other brain states, in space and time.

## 6 Methods

### 6.1 Functional Connectivity Construction

To construct the functional brain network for each subject, task, and frequency band, we calculate the Pearson’s correlation coefficient between pairs of ROIs. The coefficient measures the linear relationship between two variables and ranges from *−*1 to +1. The two variables are said to be positively correlated if there is a similar change in their values, and negatively correlated if there is an opposite change [35]. For the time series of two EEG channels *x*(*t*) and *y*(*t*) of length n, the correlation coefficient *p*_*xy*_ is given by:

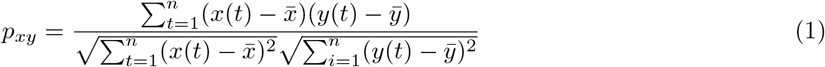

where *x*(*t*) and *y*(*t*) denote the instantaneous voltage amplitudes at instant *t*, 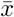 and 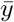 are the mean amplitudes over the entire time window, and *n* is the total number of sample points in the EEG epoch. This standardized measure of linear neuro-physiological correlation yields a symmetric 101×101 functional connectivity matrix for each experimental condition.

### 6.2 Network efficiency, global clustering and modularity and degree-preserving random networks

To characterize the global topology of the functional networks, we computed measures of network integration, segregation, and modularity. Global Efficiency (*E*), which quantifies the efficiency of parallel information flow across the network [36], is defined as the average inverse shortest path length:

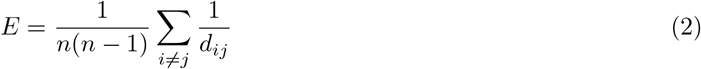

where *n* is the total number of ROIs and *d*_*ij*_ is the shortest topological distance between nodes *i* and *j*. We calculate shortest topological path between the all pair of ROIs using Networkx [37].

To measure the tendency of nodes in a graph/network to cluster together, we use the Average Clustering Coefficient(C) as a quantifying measure of segregation[25]:

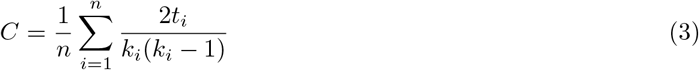

where *t*_*i*_ represents the number of triangles connected to node *i* (i.e., existing edges between the neighbors of node *i*), and *k*_*i*_ is the node’s degree [37]. Furthermore, we generate degree-preserving randomized versions of the empirical networks to compare their properties while preserving their degree distributions, which affect the topology.(The NetworkX library’s `double_edge_swap` method is utilized to randomize the graph topology while preserving the degree of the nodes[38][37]) The Newman algorithm[39] in NetworkX for Python is used to calculate the network’s Modularity (*Q*). This measure quantifies the degree to which a given network can be subdivided into multiple clusters or modules[29]:

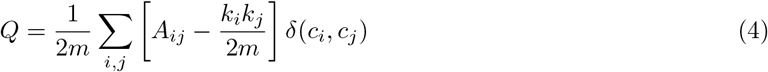

where *A* is the adjacency matrix, *k*_*i*_ = ∑_*j*_ *A*_*ij*_ is the degree of *i*^*th*^ ROI, *m* is the total number of edges in FC, and the Kronecker delta *δ*(*c*_*i*_, *c*_*j*_) equals 1 if the ROIs belong to the same assigned module (say *C*_*i*_), and 0 otherwise [37].

### 6.3 Bimodal Coefficient

We use Sarle’s Bimodal Coefficient[31] as a statistical measure to determine whether the data distribution is unimodal or bimodal. It depends on the sample size, skewness and kurtosis and is defined as follows:

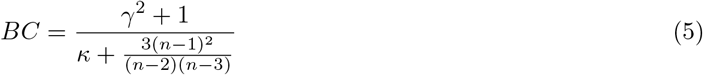

Where, *γ, κ* represents the sample skewness, and sample excess kurtosis, respectively. *n* represents the total sample size. BC value above the critical value of 0.555 is considered bimodal, and one below is unimodal.

### 6.4 Hierarchical Decomposition via *k*-core Percolation

K-core percolation involves continuously pruning ROIs with a degree less than k, starting with *k* = 1. K-Core decomposition algorithm from NetworkX is used [40]. The iterative process would continue until the minimal set of connected ROIs is reached, and further removal of ROIs based on degree would cause all ROIs to fragment. The last core of nodes is a *K*_max_-core, in which the ROIs have the maximum possible degree. [41]. The layers that are peeled off are k-shells, where the nodes in it have degree exactly *k*. Each node in the network is assigned a core number, representing the highest-order core to which it belongs. The *K*_max_ core contains the ROIs that form densely interconnected functional hubs and can be studied as a measure of networks’ robustness and topological depth [42].

### 6.5 Module Connectivity Ratio (*r*)

To further mathematically formalize the distinction between modular and anti-modular networks, we calculate the integration ratio (*r*), which quantifies the connectivity between (inter-) and within (intra-) modules. The *r* is the ratio of the probability of inter-module connections (*p*_*inter*_) to intra-module connections (*p*_*intra*_) and reflects the dominance of inter/intra-module connectivity [16].

If *n*_*i*_ and *n*_*j*_ are the number of nodes in the module *i* and *j*, and let 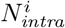 be the number of actual edges in *i*, and 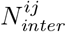 be the number of observed bridging edges connecting region *i* to region *j*. The intra-regional probability within a single module *i* is defined by the ratio of existing internal edges to the maximum possible undirected edges:

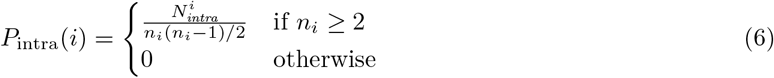

Similarly, the inter-regional probability between two distinct modules *i* and *j* is defined by the number of bridging edges divided by the maximum possible theoretical bridges between the two node sets:

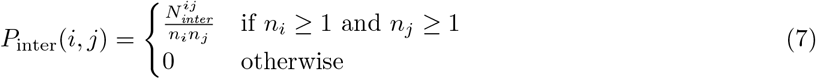

The global intra-module probability, representing the fraction of the network’s structural capacity dedicated to localized processing, is calculated as:

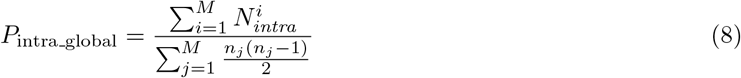

Conversely, the global inter-module probability, representing the fraction of the network dedicated to distributed, long-range communication, is defined as:

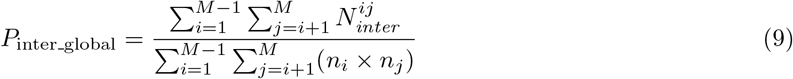

Finally, to quantify whether a specific FC is modular or anti-modular, we calculate the inter-to-intra global integration ratio (*r*), which is defined as:

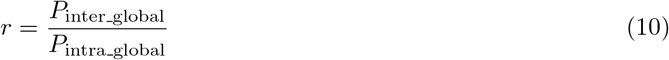

This ratio serves as a strict structural classifier: if *r >* 1, but finite: the network is partially multipartite; if *r <* 1, it is modular; and if *r≈* 1, the network exhibits no preferential modular or multipartite structure, which is characteristic of a random network topology.

## 7 Acknowledgments

We want to express our gratitude to Lakshana Balaji and Mithanshu Sukhwani for their contributions at the initial stages of this work. A.S. and S.D. thank the Department of Science and Technology (DST), Government of India, for financial support through DST INSPIRE Faculty Grant No. IFA-21-PH-276. Additionally, we would like to acknowledge IISER Tirupati for providing the High-Performance Computing facility.

## 8 Supplementary Material

### 8.1 Module connectivity ratio (r) and the Multipartite Nature of FCs

The signs of the modularity values confirm the presence of modules or anti-modules in the network, with the modules identified as the TMs. The *r* values further confirm this (Fig.S1) for *G*_+_ and *G*_*−*_ across all studied subjects, frequency bands, and states. The *r* value is more useful, as *r >* 1, but having a finite value indicates the presence of intra-TM connections. This further confirms a partial multipartite structure, not a multipartite structure, which lacks any intra-TM connections. We observe that *r* values for *G*_*−*_ networks exceed 1 for all subjects across all frequency bands and states, but can be infinite for some subjects in different bands and states. The resting state in the low frequency bands exhibits a finite *r* value for a very few subjects, and the median of the finite value is higher than the task states (Fig.S1(a)). Demonstrating the existence of more partial multipartite structure in task states, with very few subjects showing a modular or random network topology in the gamma and delta bands. Besides, the median value of *r* for the *G*_*−*_ networks across subjects is lower than the *r* values for the innermost core of these networks, exhibiting a stronger anti-modularity for the innermost core than the entire *G*_*−*_ network (Fig.S1(b)).

Moreover, Fig. S2 demonstrates the frequency with which subjects show the existence of Multipartite structures of TMs (zero intra-TM connections) in *G*_*−*_, *G*_+_, and their innermost core networks. The resting-state low-frequency band shows that most subjects (more than 17 out of 27) exhibit a multipartite structure among TMs. This number decreases in the high-frequency bands, with the gamma band showing no subjects without intra-TM connections in the resting state. The task state, on the other hand, shows fewer subjects with *r* =*∞* in low-frequency bands, with this decrease continuing into high-frequency bands, with no more than 4 subjects displaying multipartite TMs for any task state. This shows that neither task states nor resting states require negative correlations between ROIs within the same TMs in high-frequency bands. However, both rest and task states show negative correlations between ROIs within the same TMs in low-frequency bands, with the resting state showing more negative correlations within TMs than task states. This observation suggests that, in the high-frequency bands, all the ROIs within a TM do not fire simultaneously but may exhibit a lag, indicating that they engage a within-TM top-down process. Therefore, TMs can be further divided into modules to enhance anti-modularity. In contrast, in the low-frequency bands of the resting state, the TMs lack the extra layer of topological complexity. Among the task states, visual and semantic decision-making tasks exhibit the multipartite nature in more subjects than the preferential food choice task (Fig. S2 (a)). Furthermore, the innermost networks of *G*_*−*_, across all frequency bands, the resting state exhibits the absence of the negative intra-TM connections in almost all subjects, except the gamma band (Fig. S2 (b)). However, in the task state, we find the absence of the negative correlation between innermost core ROIs belonging to the same TM in most of the subjects across all the frequency bands (Fig. S2 (b)).

**Table S1:** Distribution of EEG nodes across brain cortical regions.

| Brain Region | Node Count | Nodes |
| --- | --- | --- |
| Frontal | 21 | E2, E3, E4, E5, E9, E10, E11, E12, E15, E16, E18, E19, E22, E23, E24, E26, E27, E33, E122, E123, E124 |
| Central | 22 | E6, E7, E13, E20, E28, E29, E30, E34, E35, E36, E41, E103, E104, E105, E106, E110, E111, E112, E116, E117, E118, E129 |
| Temporal | 14 | E39, E40, E44, E45, E46, E50, E57, E100, E101, E102, E108, E109, E114, E115 |
| Parietal | 20 | E31, E37, E42, E47, E51, E52, E53, E54, E55, E61, E62, E78, E79, E80, E86, E87, E92, E93, E97, E98 |
| Occipital | 24 | E58, E59, E60, E64, E65, E66, E67, E69, E70, E71, E72, E74, E75, E76, E77, E82, E83, E84, E85, E89, E90, E91, E95, E96 |

**Figure S1:**
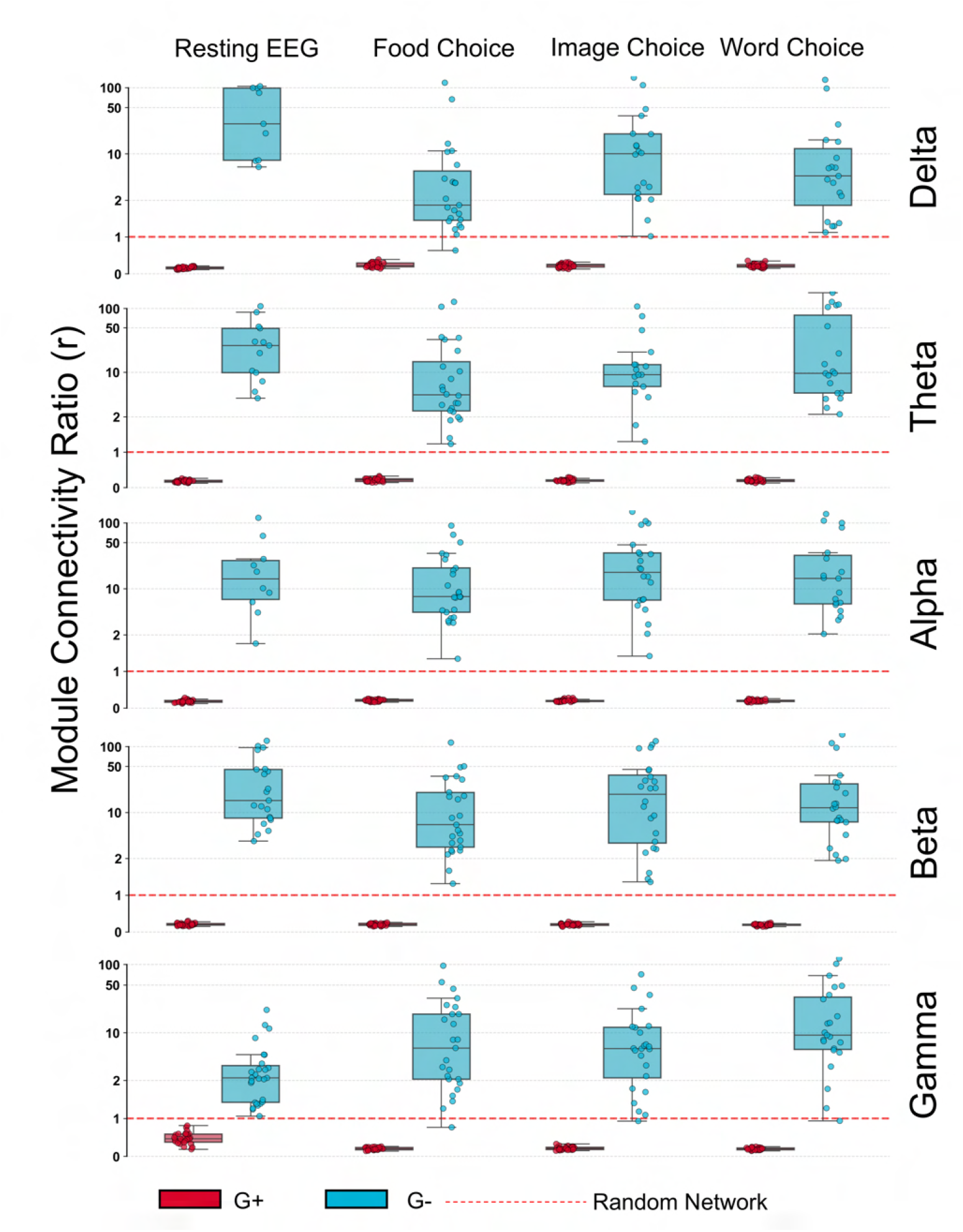
Subplots (a) and (b) illustrate the module connectivity ratios (r) for FCs of all 27 subjects across different frequency bands and states.

### 8.2 Pipeline validation

To validate the pipeline proposed in the manuscript, we run a new dataset of eye-opening rest state in the study of anxiety [28] into our pipeline and deduce the results as shown in Fig.S3(a),(b). As expected, the positively correlated network shows a modular structure, whereas the negatively correlated network shows anti-modularity, as confirmed by the fixed modularity and ‘r’ values. Further these results are compared with the original dataset used for this study in Fig.S3(c),(d). This demonstrates that anti-modularity(*Q*_*fixed*_ *<* 0) is evident in the eyes-open resting state of the subjects in *G*_*−*_ networks, irrespective of data origin.

**Figure S2:**
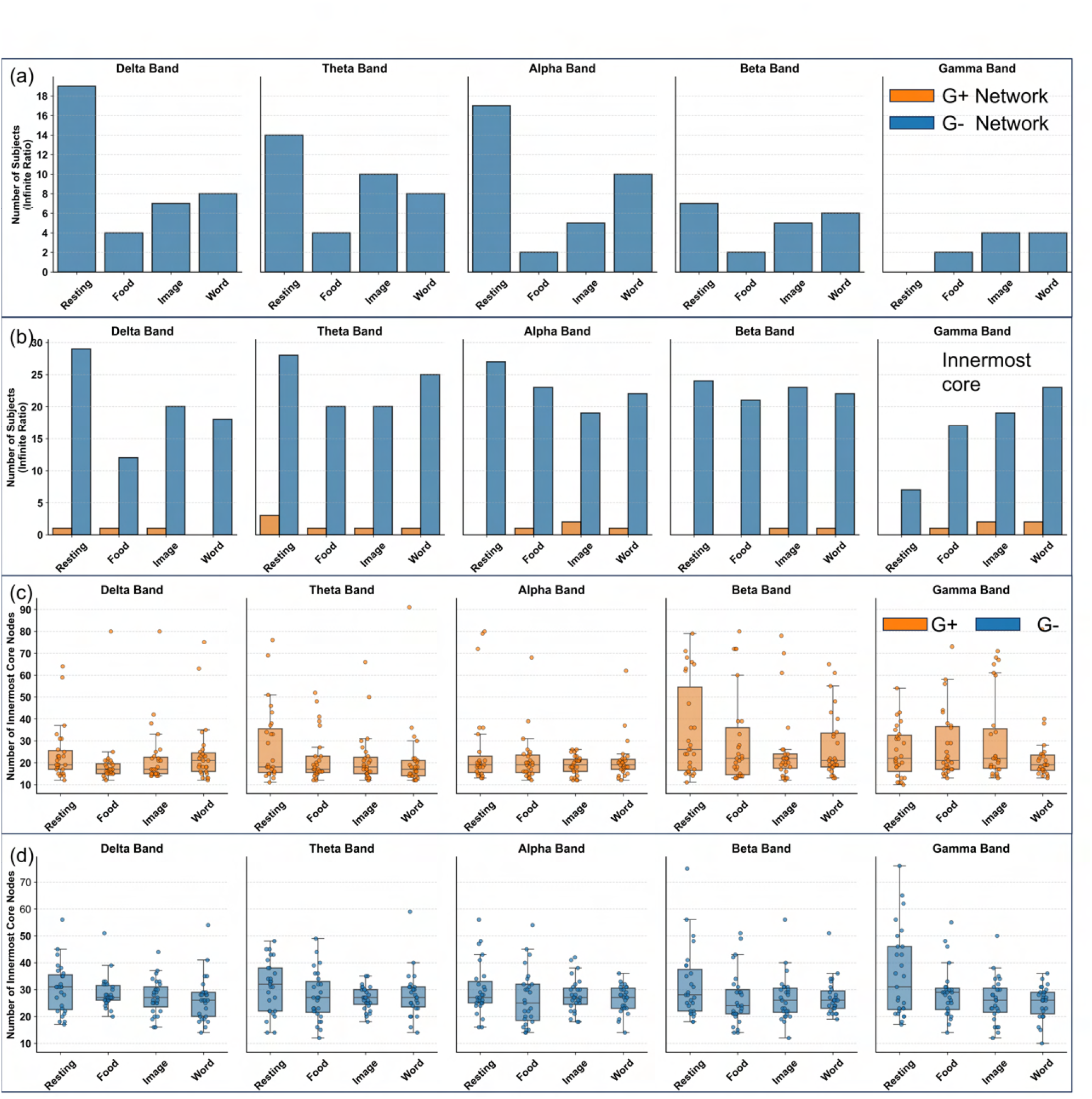
Subplots (a) and (b) plot the number of subjects with *r* = *∞*, indicating multipartite structure in the *G*_*−*_ networks and their innermost core, respectively. Box plots in (c) and (d) display the number of innermost core ROIs across subjects in *G*_+_ and *G*_*−*_ networks, respectively.

### 8.3 Edge density across TMs

When we examine the statistics on connections in Fig. S4, both within and between modules across all subjects, we observe the trend discussed below. We fix our focus on the gamma band. In *G*_+_ networks, where modularity is observed, the intra-module connections in the resting state are higher in parietal, followed by occipital and frontal modules. The central and Temporal modules show fewer edges formed within themselves. While in task states, occipital-occipital connections peaked significantly more than in the others, highlighting the module’s importance for cognitive task performance. The frontal, central, and Parietal lobes have almost identical median values, with the temporal lobe last. Also, the number of possible connections within a module should be considered, as the temporal module has 14 ROIs, as per Table S1. Overall, the task states have seen a significant increase in intra-modular connections in the correlated (+) network, with the greatest enhancement in the occipital region. On the other hand, the inter-module connections show overall suppression, which is more significant for the F-O and C-O, whereas C-T connections are increased. However, even in this case, we observe behavior in task states. F-C, C-P, and P-O are the combinations with the most connections in all task states. Note that the inter-module connections’ overall median value never crossed 0.2, which is the minimum observed in intra-modular connections, again specifying that *G*_+_ networks are highly modular and inter-modular connections are less. Overall, favored intra-modular connections in the task states are between closely neighbored modules in the (+) correlated networks.

**Figure S3:**
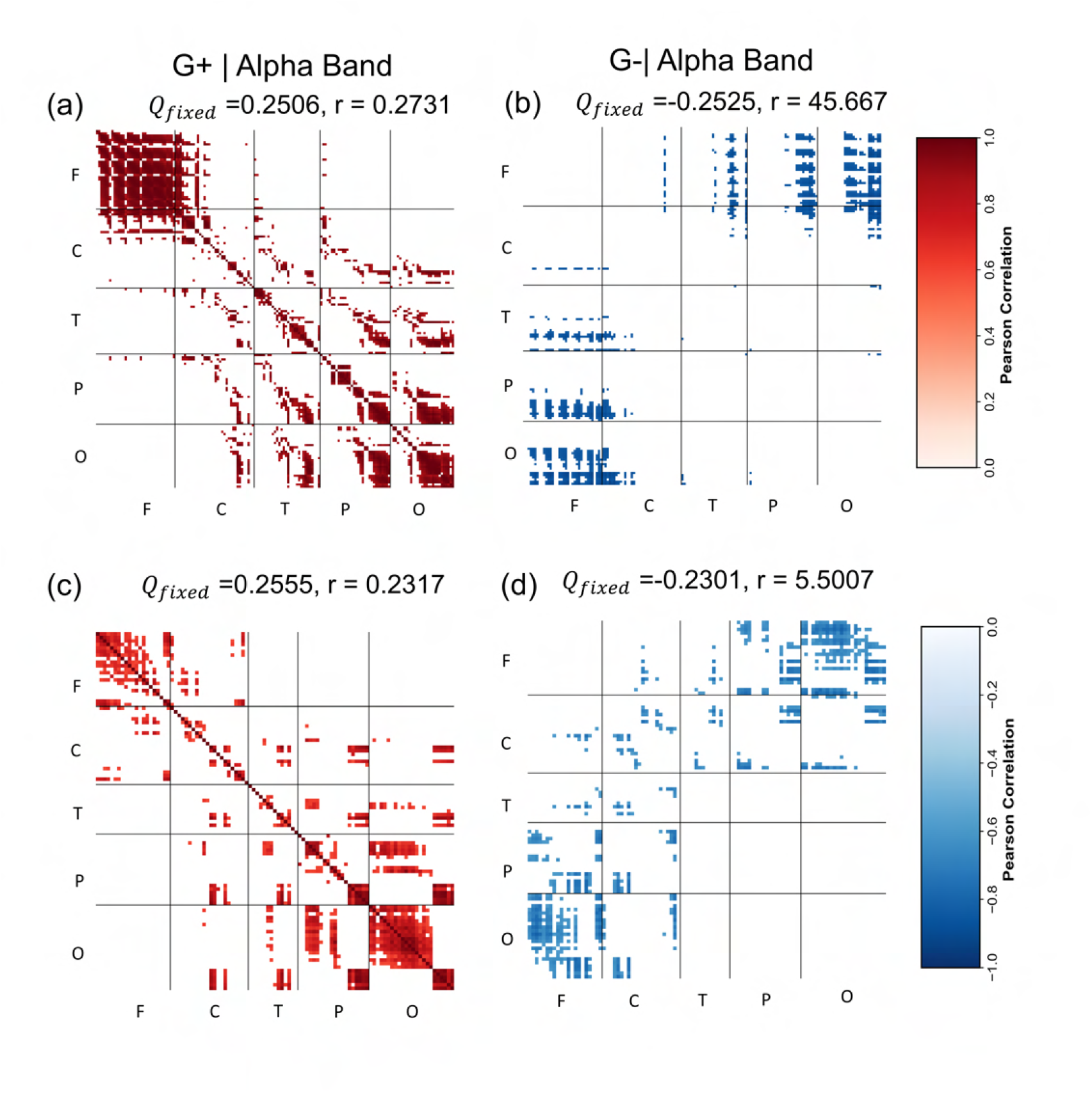
Comparing the anti-modularity in different datasets of resting state alpha band EEG networks with subjects’ eyes open. (a),(b)– Heatmaps of Pearson’s correlation in *G*_+_ and *G*_*−*_ networks constructed with density threshold as discussed in the pipeline for Sub-1032 from the dataset [28]. Five TMs are fixed, modularity (Q), and Module connectivity ratio (r) are provided for each. (c),(d)– Similarly generated heatmaps for a subject in the resting state from the dataset (Sub-07 of [43]).

In the *G*_*−*_ networks, the scenario is completely reversed. The already-proven anti-modularity is evident in the connections between modules compared to within-module connections. In the resting state, F-O and F-P are well connected, whereas in the task, F-P connections decrease, with a slight increase in F-O connections. The connectivity of C-O also enhances in task states, whereas F-C, T-P, and P-O connections decrease significantly (Figs. S4). A decrease in the negative correlations among the F-C, T-P, and P-O, along with a slight increase in positive correlations among them, indicates the need for more in-phase activity of these regions in the task states for integrating working memory with motor movement and processing visual inputs. Motor movements are in the task states to click on a choice, and the observed enhancement of positive connections within C and between F and C could explain the underlying mechanism.

**Figure S4:**
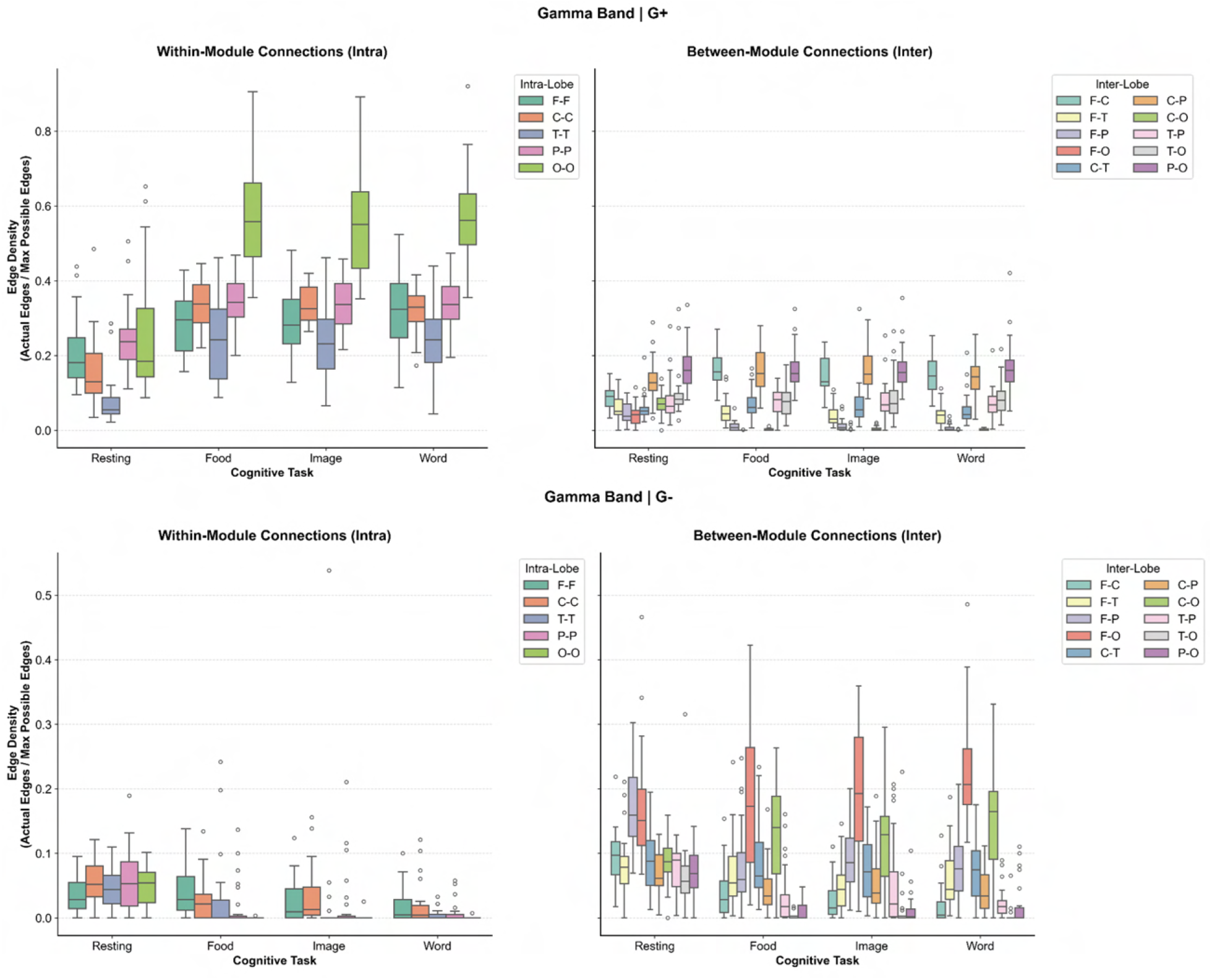
The connections/edges density for within and between modules (TMs) in the gamma band for all subjects across all the states.

### 8.4 Spread of Innermost ROIs in TMs

We provide pie charts (Fig.S5) showing the distribution of innermost ROIs across all TMs by band. The low-frequency delta and theta bands show greater posterior contributions in the resting state and a decrease in task states within *G*_+_ networks. The behavior of the Beta band is similar to that of the Gamma band, where the dominance is of the posterior. In the *G*_*−*_ networks, a balanced scenario or slightly posterior dominance exists across all tasks and bands. The ratio of frontal to occipital ROI counts indicates a more balanced structure in *G*_*−*_ networks across all bands and tasks, except in the food-choice Delta band. However, when all *G*_+_ networks showed greater heterogeneity, with more subjects’ ROIs from frontal TM across all tasks, the resting state shows a different pattern, with a greater occipital contribution. (Fig.S6).

**Figure S5:**
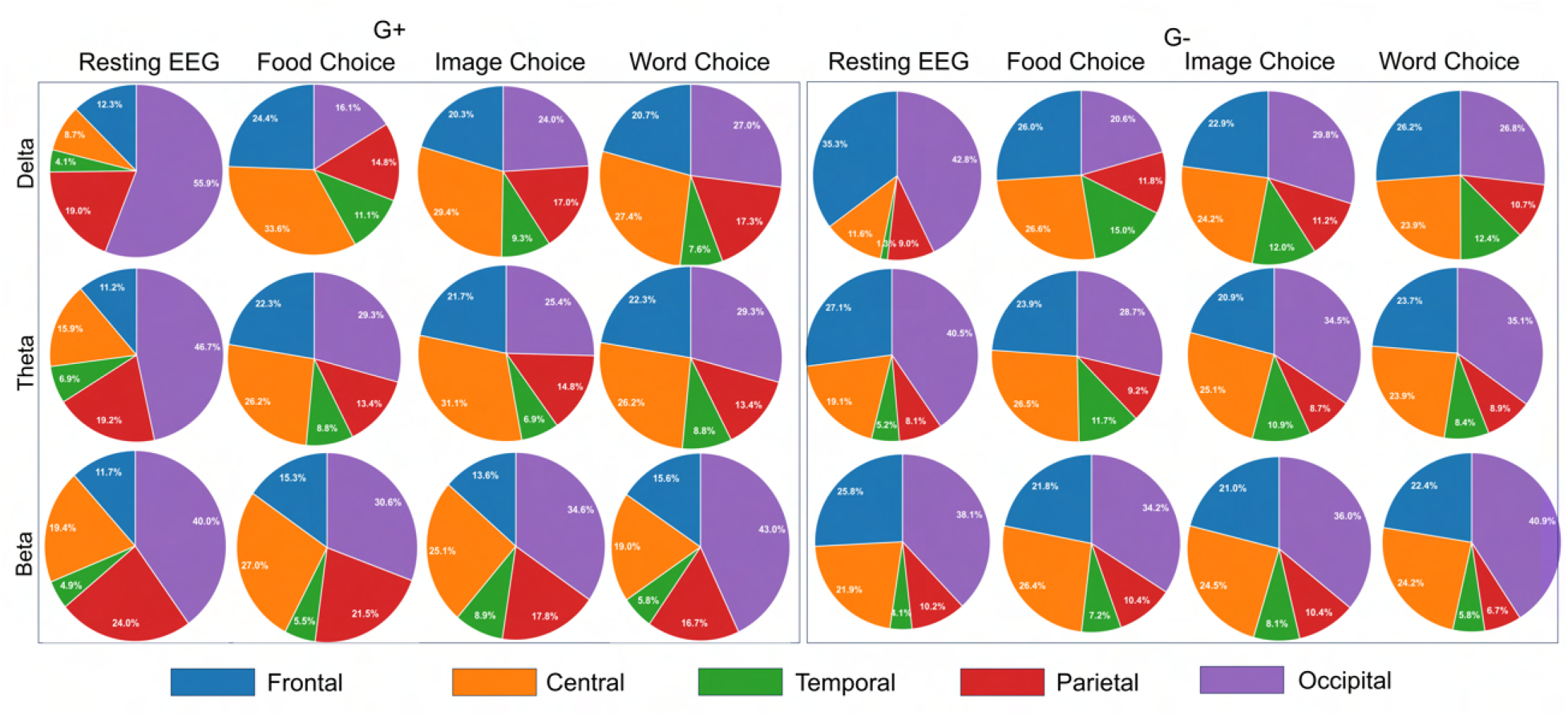
The contribution for innermost ROIs from each TM in different bands

**Figure S6:**
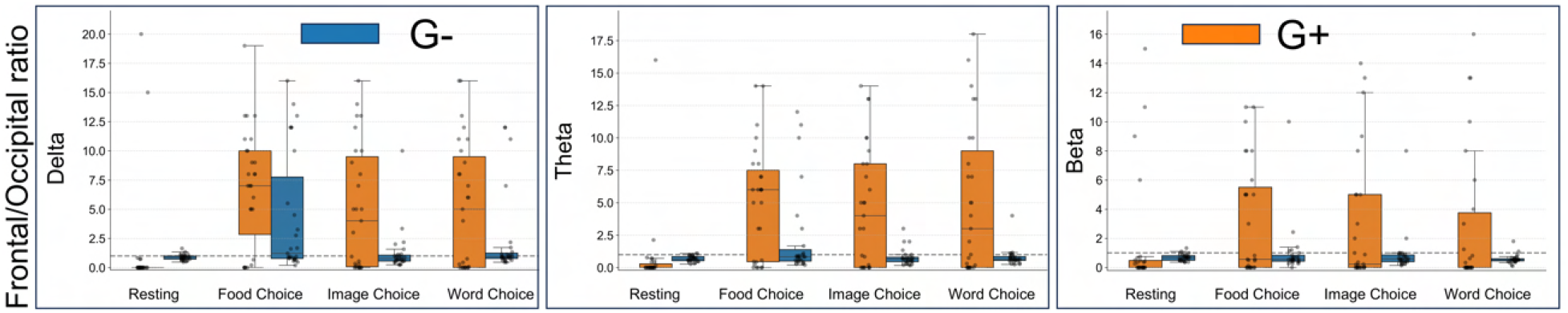
The ratio of innermost ROIs of Frontal and Occipital regions in delta, theta and beta bands.

